# Deterioration of the human transcriptome with age due to increasing intron retention and spurious splicing

**DOI:** 10.1101/2022.03.14.484341

**Authors:** Marco Mariotti, Csaba Kerepesi, Winona Oliveros, Marta Mele, Vadim N. Gladyshev

## Abstract

Adult aging is characterized by a progressive deterioration of biological functions at physiological, cellular and molecular levels, but its damaging effects on the transcriptome are not well characterized. Here, by analyzing splicing patterns in ∼1,000 human subjects sampled across multiple tissues, we found that splicing fidelity declines with age. Most prominently, genuine introns fail to be spliced out, manifesting as a broad surge in intron retention, and this is exacerbated by the increase in diverse spurious exon-exon junctions with age. Both of these effects are prominently detected in the majority of human tissues. Collectively, they result in the progressive deterioration of the active transcriptome, wherein functional mRNAs are increasingly diluted with non-functional splicing isoforms. We discuss the concept of “splicing damage” and formulate methods to quantify it. Using these tools, we show that splicing damage increases both with age and with the incidence of diseases. Altogether, this work uncovers transcriptome damage as a critical molecular indicator of human aging and healthspan.

## Introduction

Aging is accompanied by a vast array of molecular, cellular and physiological changes, generally characterized by a global decline in biological function and health throughout adulthood ^1^. By applying diverse molecular profiling techniques, researchers have now begun elucidating the epigenetic, transcriptional, and metabolic changes that accompany aging ^2–5^. Splicing is an important layer of gene expression leading to gene product diversification and regulation.

In the last ten years, several studies reported changes in alternative splicing with age, as reviewed in ^6^. A 2011 study profiled gene expression in leukocytes by microarray analysis and found that genes involved in mRNA processing and splicing significantly changed with age, leading authors to hypothesize that splicing patterns may also be altered ^7^. This view was further reinforced with the advent of high-throughput RNA sequencing. A multi-tissue study of mouse showed that alternative splicing isoform usage increased with age, and splicing machinery genes were themselves affected ^8^. Both these observations were confirmed by a recent multi-tissue study in human, which identified as many as ∼50,000 splicing events whose frequency correlated with age, mostly with tissue-specific patterns ^9^.

Several studies analyzed splicing changes occurring specifically in the aging brain to investigate potential mechanisms of cognitive disease, in particular Alzheimer’s disease (AD) ^10–15^. From this research, one type of splicing event is emerging as relevant for aging and disease: intron retention (IR), in which an intron is retained in the final mRNA instead of being spliced out (Fig. 1a). It has been observed that, in the human brain, many introns show increased levels of retention with age ^11^. The same effect was seen in brains of patients with AD compared to controls ^14,15^. Notably, an age-dependent increase in many IR events has been shown also in rhesus macaque brains ^11^, mouse brains ^15^, fruit fly heads ^15^, and *Caenorhabditis elegans* ^16,17^. These observations have put IR research in focus, but much remain to be understood. In particular, a global study of IR across genes and tissues is lacking, which could address whether IR represents a general signature of human aging.

**Fig. 1:**
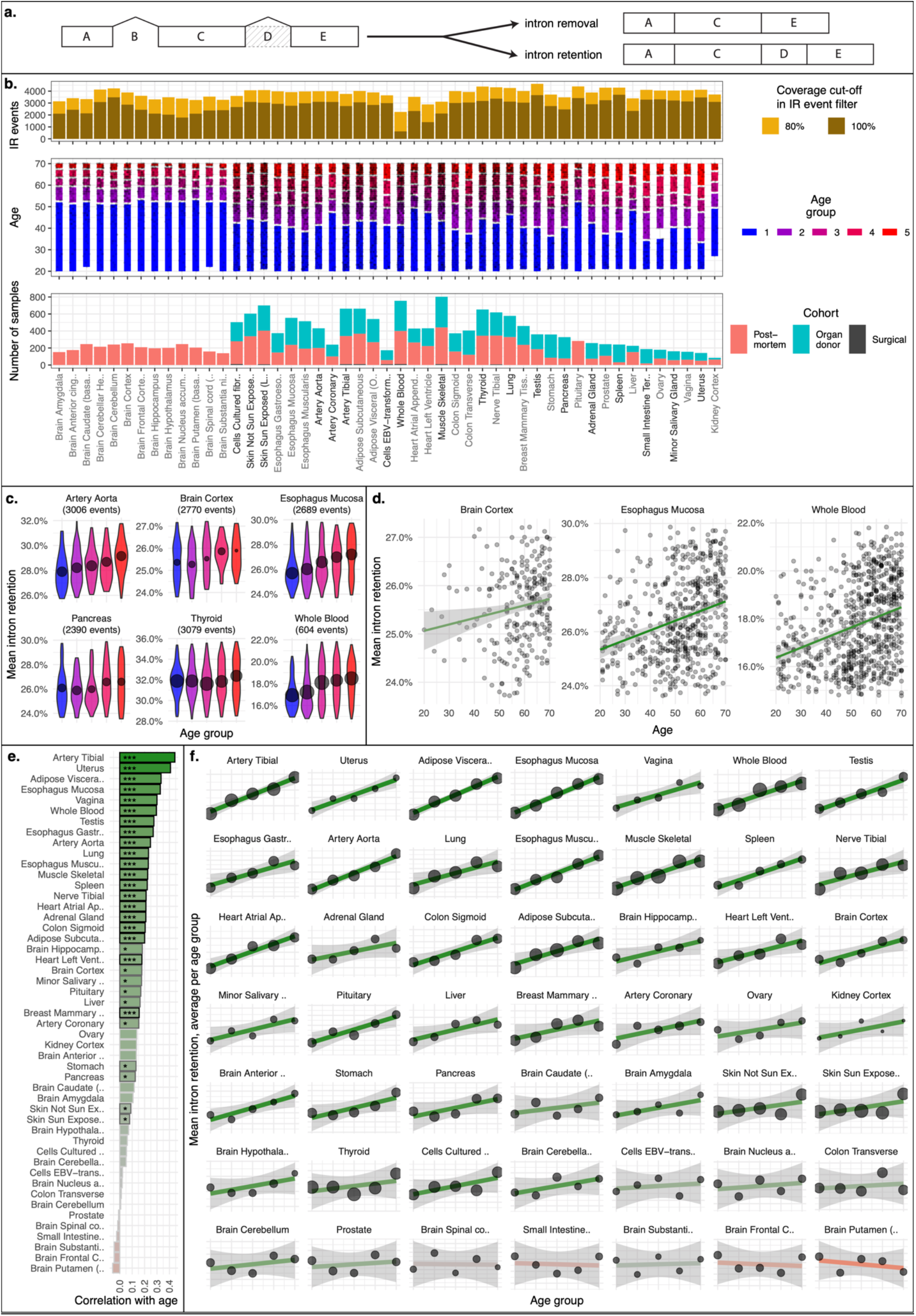
Change with age of mean IR in GTEx samples. (a) Schematic representation of an IR event, showing the mRNA resulting from either complete removal or retention of the intron marked as “D”. (b) Data overview showing the number of RNAseq samples per subtissue colored by GTEx cohort; the age distribution and age group segmentation; and the number of intron retention (IR) events passing the coverage filter, colored by cut-off (see Methods). Subtissues are grouped by tissue, as depicted by the alternating grey and black text color of x-axis labels, and ordered by the number of samples per tissue. For subsequent analyses, the sets of IR events passing the coverage cut-off of 100% were used, unless otherwise noted. (c) Distribution of mean intron retention per sample (average PSI value of IR events), split by age groups, for six representative subtissues. Black points indicate mean values per age group, with the size proportional to the number of samples. (d) Raw values and trend lines for three representative subtissues. (e) Pearson correlation coefficients between IR burden and age per subtissue. Stars indicate statistical significance (***= FDR < 0.05; *= nominal p-value < 0.05). (f) Summary of age changes of mean IR for all subtissues, showing average values per age group (analogously to panel (c)) with trend lines. Colors in panels (d-f) represent correlation coefficients (green= positive, red= negative). In panels (c) and (d), outliers (top and bottom 3%) were removed for visualization purposes.

In this work, we analyzed IR changes with age in the largest human transcriptomic dataset available, GTEx, encompassing approximately a thousand individuals sampled across multiple tissues. We discovered a systematic age-related increase of the transcriptome-wide level of IR across the great majority of tissues. We observed that the age-associated trend of individual introns is mostly consistent across tissues and relates to the functional consequences of each IR event. Indeed, the emerging picture is of an age-dependent drift from a “healthy” transcriptome, as functional transcripts are increasingly diluted by non-functional splicing variants. This predominantly comes from progressively higher levels of IR, but also from increasing occurrence of spurious splicing of functional exonic portions. Based on these advances, we defined new metrics to quantify these “splicing damage” effects, and analyzed their age trajectory, determinants and correlated transcriptomic changes. Altogether, we reveal splicing damage as a new indicator of deterioration of molecular function, relevant for studying and quantifying the effects of aging and disease.

## Results

### Quantification of splicing across tissues and ages

To investigate the impact of aging on intron retention (IR) in humans, we analyzed 17,329 RNAseq samples encompassing 49 subtissues of 948 subjects (Fig. 1b), constituting the GTEx dataset v8 ^18^ after filtering (see Methods). We defined age groups for visualization and analysis by splitting samples in 5 quantiles for subtissues (Fig. 1b). The age boundaries of groups varied between subtissues, mostly reflecting different cohort compositions (<u>Fig. S1</u>).

IR was quantified using the program Suppa ^19^. This program derives the set of IR events from a gene annotation, selecting all introns which are spliced in at least one annotated transcript and retained in at least another isoform of the same gene. IR events are quantified separately in each sample and expressed in the form of PSI values (percent-spliced-in) ranging from 0 (complete intron removal) to 1 (complete retention). Quantification requires transcript isoforms to be expressed. We filtered Suppa results to retain only the events quantified in abundant samples across all ages (“coverage cut-off”; Methods). For most analyses, we applied the strictest coverage cut-off: for each subtissue we considered only the IR events quantified in all samples. These sets ranged in size from 624 to 3,696 IR events across subtissues, with an average of 2,773 (Fig. 1b). These events corresponded almost entirely to introns in protein coding genes, both because most other gene types were intronless, and because coding genes exhibited higher expression levels, allowing them to pass our filtering procedure (<u>Fig. S2</u>).

### IR increases with age

First, for each sample, we computed the mean IR, defined as the average PSI value of IR events quantified in all samples of that subtissue. Note that this measure is dependent on the set of IR events considered, defined here per subtissue. Strikingly, the mean IR exhibited a robust increase with age in most subtissues (Fig. 1b-e). We observed a positive correlation in 43 of 49 subtissues (Fig. 1d), reaching nominal significance in 30 cases (Pearson correlation p-value <0.05) and FDR significance in 20 cases (Benjamini-Hochberg adjusted p-value <0.05). None of the tissues showed a significant decrease in IR with age. To assess whether the IR increase with age could be due to covariates, we fit a linear mixed model with mean IR as function of age (fixed effect) with subject, subtissue and cohort as random intercepts. In this model, age showed a significant positive slope with mean IR (p-value<0.05). We explored many additional covariates recorded in GTEx (sex, autolysis score, collection site, race, ethnicity, and Hardy scale death circumstance), but we ultimately dismissed them as they accounted for less than 1% of variance when included in the model as random effects. We also fit a model for each subtissue separately (with cohort as the only random intercept), which resulted in positive slope in 37 subtissues, with 14 of them reaching significance (p-value< 0.05). These results show that the global level of IR shows a robust age-dependent increase, detected across the remarkable diversity of GTEx samples.

Next, we investigated how individual IR events contributed to the observed increase of mean IR with age. For each subtissue and for each IR event separately, we fit a linear model (PSI ∼ age), and built a matrix with the resulting slopes, wherein each value represented the trend with age of a single IR event in a particular subtissue. We then used this matrix to cluster events and subtissues (Fig. 2a). For this analysis, we relaxed our filtering criteria, expanding IR event sets in order to better assess the consistency of individual IR events across subtissues. We used a coverage cut-off of 80% (Methods) per subtissue, and then excluded events quantified in <25% of subtissues for visualization purposes. IR events (i.e., introns) separated in roughly 4 clusters (Fig. 2a): two clusters featuring a clear IR increase with age (the “I1” cluster of 326 events with the highest slope, and the “I2” cluster of 1,195 events), one large cluster with little or no trend with age (the “F” cluster for “flat”, 1,506 events) and a small cluster which decreased with age (“D” cluster, 135 events). Although I1 included only 6% of IR events, its slopes summed up to 44% of the total (median value across subtissues; <u>Fig. S3</u>). The slope of any given IR event was typically consistent across subtissues, with one remarkable exception: cultured cells showed basically no correlation with other subtissues (Fig. 2b).

**Fig. 2.**
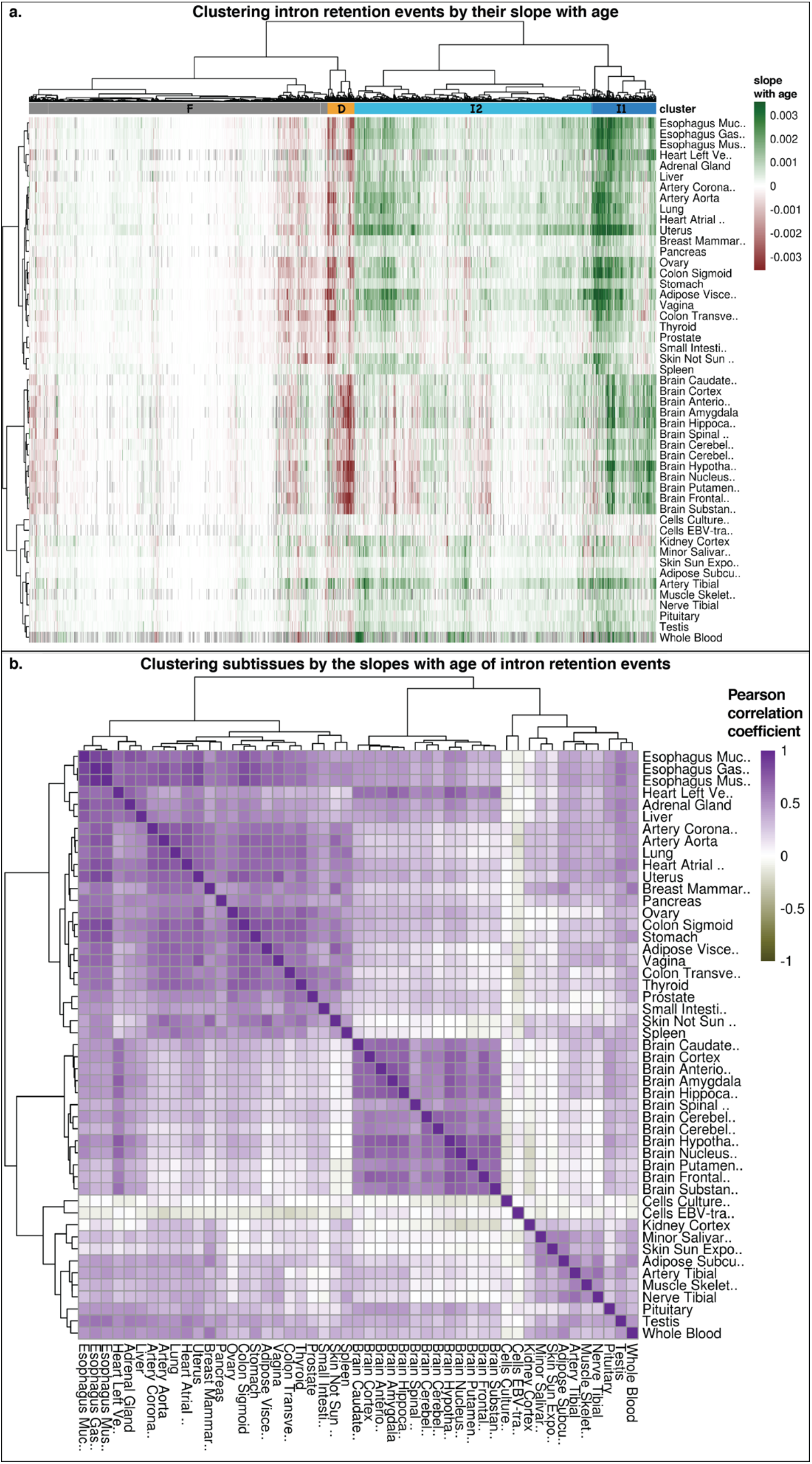
Clustering intron retention events and subtissues. (a) Clustered heatmap of IR events (columns) and subtissues (rows). Clustering was performed based on the slopes of IR with age of individual IR events. The slopes are represented here in color: green for increase, red for decrease (color saturation set at 25% of the global maximum slope). The clustering structure on top was used to define 4 clusters of IR events: F (flat, grey), D (decrease, orange), I2 (increase, light blue), I1 (strong increase, dark blue). (b) Clustered heatmap of subtissues based on the trend with age of IR events. Color designates the Pearson correlation coefficient between pairs of subtissues, while the clustering structure of subtissues is taken from panel (a).

### Non-functional splicing isoforms increase with age

Depending on intron location within the transcript, IR may have very diverse consequences on the synthesis of protein products. For example, the retention of an intron located between two coding exons alters the encoded protein sequence, typically in a dramatic fashion (through frameshifts or premature stop codons). On the other hand, the retention of an intron in the 3’UTR does not change the protein. Importantly, to assess the functional consequences of IR events, one needs to define directionality, i.e., choose a reference mRNA isoform per gene. Intuitively, we think of IR as an alteration wherein an extra intronic sequence is included in the final mRNA. Yet, while most IR events fit this intuition, others involve introns which are spliced out only in secondary isoforms. For these, it is more natural to think of intron retention as the normal status, and its exclusion (i.e., intron removal) as the alteration. Notably, the PSI values have opposite functional interpretations in the two cases, so that we must split events into categories for a meaningful analysis.

In view of this, we selected a reference transcript for each gene, taking the “principal” functional isoform as defined by APPRIS ^20^. When several were present, we selected the top expressed one per subtissue. Then, we assessed the kind of alterations each IR event constituted when compared to that isoform. We defined 11 possible types of consequences (Fig. 3a; detailed in Methods), composed of two principal groups: intron inclusions (“IR-in”) and intron exclusions (“IR-ex”). Inclusions correspond to the retention of introns that are normally spliced out to produce the reference mRNA, i.e., the typical intuition of IR. They can introduce frameshifts (“in-FS” category) or premature stop codons (“in-PTC”), or lead to other consequences (“in-UTR” or “in-pep”). Exclusions represent the removal by splicing of portions of exons present in the reference mRNA, resulting in frameshifts (“ex-FS”) or other alterations (“ex-Nter”, “ex-Cter”, “ex-UTR”, “ex-pep”). Other functional categories also exist (“noncoding” and “other”; Methods). Note the opposite interpretations of PSI values: for inclusions, the functional isoform is produced upon intron removal (PSI=0), while for exclusions, it is produced upon its retention (PSI=1). This is apparent in the different distributions of mean PSI of inclusions and exclusions (Fig. 3b), which reflect their natural functional state.

**Fig. 3.**
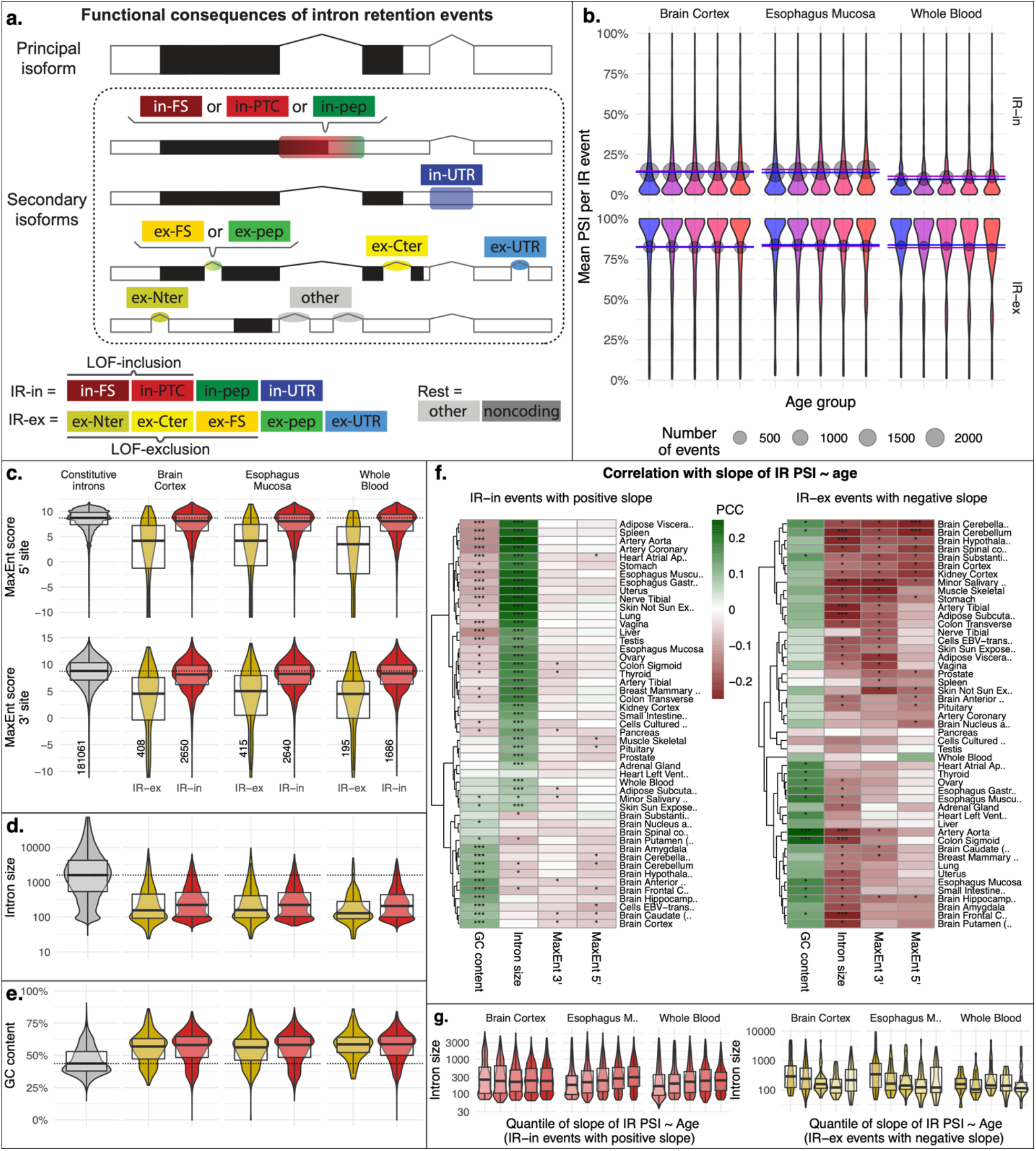
Functional consequences of IR events and features affecting their age-related change. (a) Examples of the 11 types of functional consequences attributed to each IR event (Methods). Categories grouping together multiple consequences are shown at the bottom. In the diagrams, coding sequences are colored in black. (b) Distribution of the mean PSI values of inclusion (IR-in) and exclusion (IR-ex) events in three representative subtissues. The mean PSI is computed here per event, averaging across samples, separately for each subtissue and age group. Black points indicate the average of mean values per age group per category (IR-in or IR-ex), with the size proportional to the number of events. Colored horizontal lines correspond to the average in the youngest and oldest age groups. (c) Comparison of the distribution of splicing site strength (MaxEnt score) between constitutive introns (i.e., always spliced out; Methods), IR-ex events, and IR-in events. Three representative subtissues are shown. (d) and (e) The same comparison of (c) for intron size and GC content, respectively. The number of events in each category is indicated at the bottom of panel (c). (f) Pearson correlation coefficient of each of these intron features with the slopes per event of the regression PSI ∼ age, in the form of a clustered heatmap. Stars indicate statistical significance (***= FDR < 0.05; *= nominal p-value < 0.05). (g) Distribution of intron size, splitting IR events in quantiles by their slope (see <u>Fig. S5</u> for other subtissues and other features). Note that for age-increasing IR-in events ((g) left), the fifth quantile corresponds to the largest increase, while for age-decreasing IR-ex ((g) right), the first quantile corresponds to the largest decrease.

To focus on damaging effects, we further defined groups of events expected to result in loss-of-function (LOF), leading to the production of non-functional transcripts instead of functional reference isoforms. We defined LOF-inclusions (comprising of “in-FS” and “in-PTC” events) and LOF-exclusions (including “ex-Nter”, “ex-Cter”, and “ex-FS”). Strikingly, we noticed that LOF-inclusions and LOF-exclusions followed opposite trends with age. The “D” cluster of IR events (whose PSI decreases with age) contained more LOF-exclusions events than other clusters (Fisher exact tests, Benjamini-Hochberg adjusted p-values < 2.03e-3 in every subtissue); and the “I1” and “I2” clusters (whose PSI increases with age) contained more LOF-inclusions than the other clusters (adjusted p-values < 7.36e-15 in every subtissue) (<u>Fig. S4</u>a). Indeed, the slope with age was on average positive for LOF-inclusions and negative for LOF-exclusions for the vast majority of subtissues (<u>Fig. S4</u>b), indicating that, in older ages, both classes drift away from the functional state. Altogether, our results indicate that splicing fidelity progressively deteriorates with age, both by failing to splice out genuine introns (“under-splicing”) and by splicing out spurious introns (“over-splicing”), thus impairing the production of functional mRNAs.

### Introns with altered occurrence with age have specific sequence features

Next, we set to identify sequence features characterizing the IR trend with age for each intron. Previous research pointed to the strength of splice sites, the length of introns, and their GC content as relevant attributes ^21,22^. We compared these features among three groups of introns: those with inclusions as functional consequence (“IR-in”, Fig. 3a), those with exclusions instead (“IR-ex”), and a set of constitutively spliced out introns (Methods). We predicted the strength of the 5’ and 3’ splice sites of each intron using the MaxEnt program ^23^. Our results (Fig. 3c) show that the splice sites of IR-ex introns exhibited markedly lower scores than both IR-in and constitutive introns (Wilcoxon–Mann–Whitney tests, p-value<1e-38 for all comparisons in every subtissue), consistent with the idea that IR-ex are non-functional splicing events. Splice sites of IR-in introns featured overall lower scores than constitutive introns (p-value<1e-15), though their score distribution was largely overlapping. Intron length (Fig. 3d) was also on average larger in constitutive introns than in IR-in or IR-ex introns (p-value<1e-34), with the former being slightly longer than the latter (p-value<5e-3). GC content (Fig. 3e) was significantly lower in constitutive introns (p-value<1e-42), and nearly identical between IR-in and IR-ex introns (N.S.). These findings are consistent with the previously reported characteristics of retained introns ^15,21,22^.

Next, we examined whether these features were relevant for the degree of age-dependent IR trends. We computed the slopes of IR with age resulting from fitting a linear model PSI ∼ age for each event and subtissue separately. We initially focused on IR-in events (the majority), which, as already discussed, exhibited positive slopes on average. We explicitly selected those IR-in events with positive slopes and examined the correlation between slope and any of the features listed above. Most notably, intron size significantly positively correlated with the slope in most subtissues (Fig. 3f), so that introns with the highest IR increase tended to be longer (Fig. 3g; <u>Fig. S5</u>). Splice site strength negatively correlated with slope in most subtissues (Fig. 3f; <u>Fig. S5</u>), though the correlation did not reach statistical significance after multiple testing correction. The GC content showed a puzzling relationship with slope, with opposite significant effects in different subtissues (Fig. 3f; <u>Fig. S5</u>). Next, we considered the IR-ex events (which tend to decrease in PSI value with age), selected those with negative slope, and performed an analogous analysis. Splicing site strength and intron size correlated negatively with their slope, while GC content showed a positive effect (Fig. 3f-g; <u>Fig. S5</u>), so that the events with the strongest PSI decrease (corresponding to the production of “impaired” deleted isoforms) showed a tendency to be longer, AT-richer, and have stronger splice sites.

Taken together, our results indicate that genuine introns (IR-in) failing to be spliced out with age tend to have weaker splice sites and to be longer (and yet, they are shorter than constitutively spliced out introns). On the other hand, spurious introns (IR-ex) that are increasingly spliced out with age are most similar to constitutive introns, in that they tend to be longer, GC-richer, and possess stronger splice sites. We must note, however, that these effects have relatively small magnitudes and are not entirely consistent. Therefore, we conclude that these features are only a part of the story, and other unknown characteristics must contribute to determine trends with age of individual introns.

### Transcriptional signatures of IR include age-and splicing-related pathways

Seeking further insights into the mechanism of age-associated IR increase, we set out to characterize transcriptional changes correlated with IR. For context, we first analyzed the broad landscape of transcriptional changes with age in our dataset. First, we computed the relative representation of different gene types in the transcriptome, by summing up gene expression values (TPM) per gene type category (e.g., “protein-coding”, “miRNA”, etc.) in each sample, and we then analyzed the correlation with age. The transcriptome composition changed remarkably with age (Fig. 4a), with subtissues roughly split in two groups (independent of cohort). In one group, many non-coding gene types (e.g., miRNA, snRNA, lincRNA) increased their total transcriptomic output with age, while in the other subtissues, these decreased instead.

**Fig. 4.**
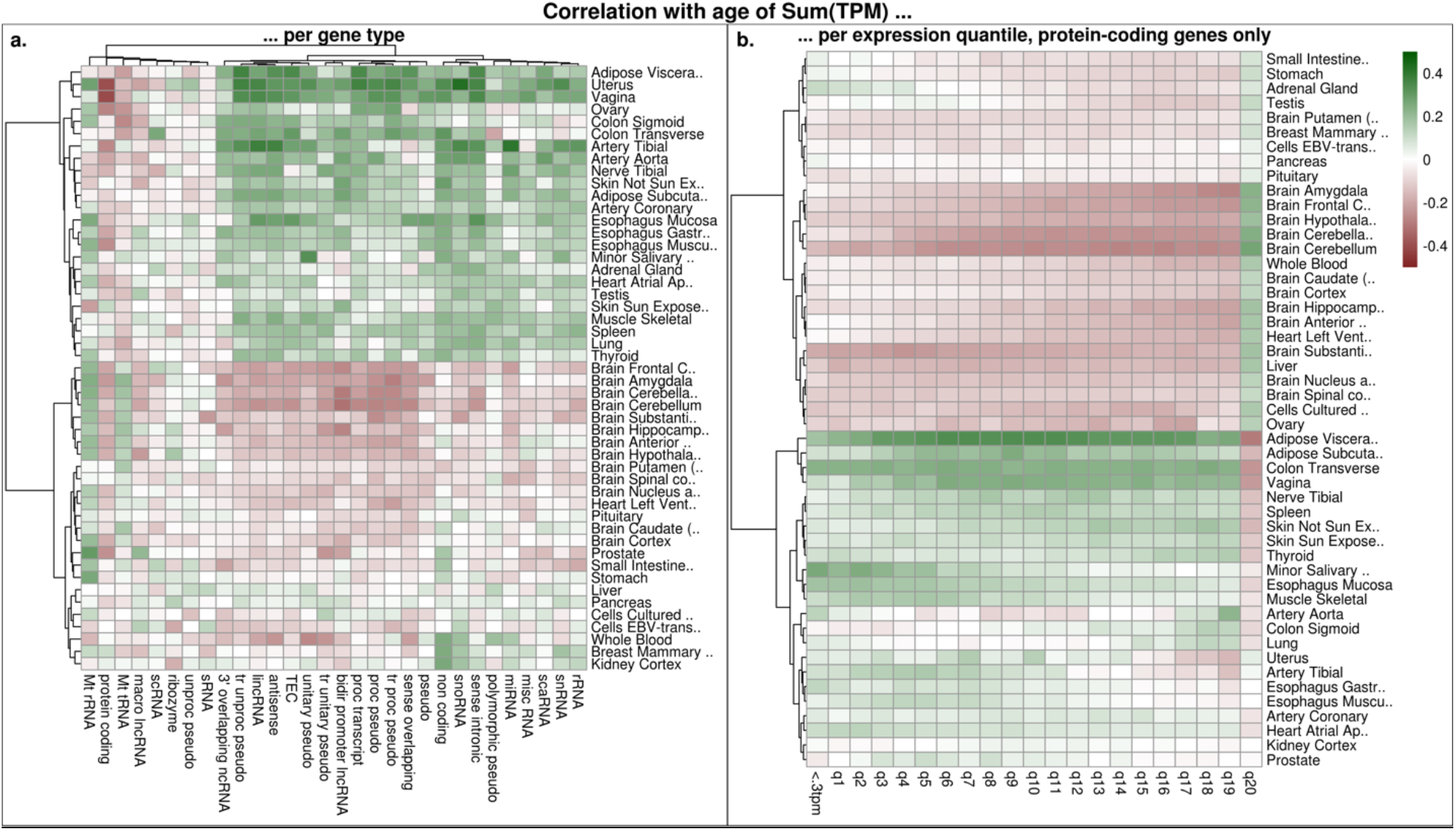
Global shift in gene expression with age. (a) Clustered heatmap showing the Pearson correlation of age with the sum of gene expression values in transcript-per-million (TPM), splitting genes by type (some are shown with abbreviated labels: tr=“transcribed”, proc=“processed”, pseudo= “pseudogene”, bidir= “bidirectional”). (b) Here, the same measure was computed after partitioning protein-coding genes in 20 quantiles, based on their mean TPM across samples. The group of genes with TPM<0.3 (omitted from other quantiles) is also shown.

We further tested whether the shape of the distribution of expression values changed with age, e.g., altering the transcriptome composition in terms of lowly abundant vs highly expressed genes. We partitioned protein coding genes in expression quantiles, computed the sum of TPM values per quantile, and examined its correlation with age (Fig. 4b). Subtissues clustered quite sharply in two groups, roughly corresponding to those in our previous analysis. In around half of the subtissues (including brain samples), the total number of transcript molecules from most expressed genes increased with age, therefore reducing transcript diversity; conversely, the relative transcriptome representation of all other gene quantiles decreased. In the other half of subtissues, (including adipose tissue and skin, among others), just the opposite occurred i.e., highly expressed genes decreased their transcript levels. Note that, since measured expression levels are relative to the total transcriptome, we cannot unequivocally differentiate the two scenarios: the observed pattern may be due to a change in transcript levels from highly expressed genes, or due to the opposite change from the rest of genes. Yet, the former is a more parsimonious explanation, and thus seems much more likely.

In our quest for transcriptional changes correlated with IR, we realized that these global shifts in transcriptome composition posed a technical challenge, since they can systematically skew the distribution of the t-value statistic used in regression analysis (<u>Fig. S6</u>). Fortunately, the adoption of methods specifically designed for gene expression modelling (Limma and edgeR) remedied this issue (<u>Fig. S6</u>). We thus used Limma to detect genes whose expression correlated with the mean IR level per sample, while including age, cohort, and sex as covariates, and analyzed gene set enrichment. We performed this analysis for each subtissue separately, then focused on those gene sets showing a consistent effect across subtissues (<u>Fig. S7</u>), which resulted in 44 pathways negatively correlated with IR (Fig. 5a). This set was redundant, since the same genes may drive enrichment of multiple pathways. Therefore, we visualized the signal components (t-values of individual genes) together with the gene set overlap among pathways (Fig. 5c-d), allowing us to pinpoint these processes: spliceosome, ribosome, protein export, proteosome, endocytosis, oxidative phosphorylation, lysosome, and several metabolic pathways.

**Fig. 5.**
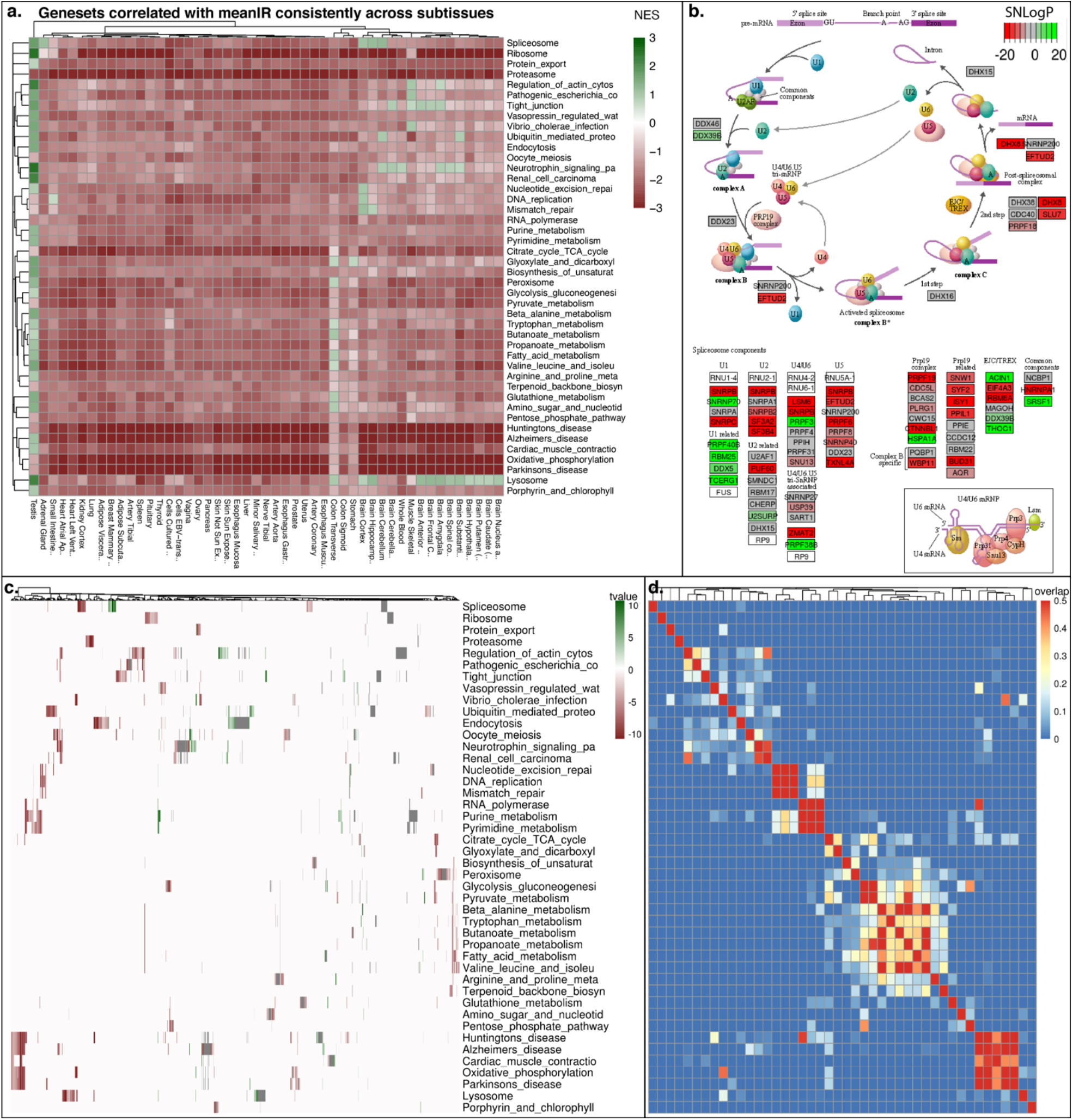
Pathways whose expression correlates with mean IR level across samples. (a) Gene sets correlated with mean IR consistently across subtissues (the full list of enriched gene sets are shown in <u>Fig. S7</u>). Mean IR is computed here per sample, as the average PSI value of IR events quantified in all samples of that subtissue. Rows represent gene sets (KEGG pathways), columns represent subtissues, and color designates the Normalized Enrichment Score (NES). (b) Results for the spliceosome in a representative subtissue (esophagus mucosa). Significant genes (p-value<0.05) are colored by their SNLogP (negative log of the p-value associated to the slope gene expression ∼ meanIR, with sign to indicate positive or negative correlation). (c) Breakdown of the gene sets in panel (a) to individual gene level: rows represent gene sets, columns represent genes (belonging to one or more sets), and color represents the associated t-value. (d) Content overlap of different gene sets. Both rows and columns represent gene sets, aligned to panel (c).

Due to their direct relevance for IR, we further analyzed spliceosome genes (Fig. 5b). Many splicing factors and related proteins, encompassing all the steps of the splicing process, showed significant negative correlation with mean IR (after accounting for covariates). Interestingly, these included the majority of proteins which form (or are related to) the Prp19 complex, a major player in splicing (<u>Fig. S8</u>).

While inspecting the list of processes correlated with IR, we noticed that several of them have been previously associated with age. Indeed, when we searched for pathways whose gene expression correlated with age in our dataset (without taking IR into account), we found again the proteosome, oxidative phosphorylation, and proteolysis as negatively correlated pathways (<u>Fig. S9</u>). Note that age was considered as a co-variate in our previous analysis model, implying that the correlation of these pathways with IR goes beyond what is explained by age alone.

Lastly, to test whether the increasing transcriptome deterioration due to under-splicing and over-splicing was linked to distinct mechanisms, we searched for pathways correlated to the mean of IR-in or the mean of IR-ex separately. Both these analyses (<u>Fig. S10</u>) resulted roughly in the same gene sets described above for mean IR.

### The usage of rare exon-exon junctions increases with age

At the onset of examining age changes, our expectation was that splicing fidelity may be decreasing with age, leading to a more diverse (and less functional) transcriptome. Our analysis of IR events described above supported this hypothesis, but with a limitation. The IR events quantified by Suppa rely on a fixed genomic annotation, and, as such, they do not fully capture transcriptome diversity generated by “splicing noise” (i.e., rare spurious splicing events), since many unannotated exon junctions may appear. Therefore, we turned to the program Leafcutter ^24^, which performs *de novo* quantification of exon-exon junctions by identifying and processing split-mapped RNAseq reads. Leafcutter cannot quantify IR, but it is well suited for the analysis of junctions. We used Leafcutter data to calculate the proportion of reads supporting rare exon-exon junctions in each GTEx sample, a measure hereafter referred to as “rare junction usage” (see <u>Fig. S11</u> for its definition). Our analysis shows that rare junction usage increases with age in the great majority of subtissues, consistent with an age-dependent deterioration of splicing fidelity. Notably, this trend was significant even in some subtissues without a strong age-dependent signal in mean IR (e.g. several brain sections) (Fig. 6). We also took an alternative approach to quantify splicing noise, calculating the proportion of Leafcutter reads supporting junctions which are missing from the annotation (“unannotated junction usage”), and results were very similar (Fig. 6).

**Fig. 6.**
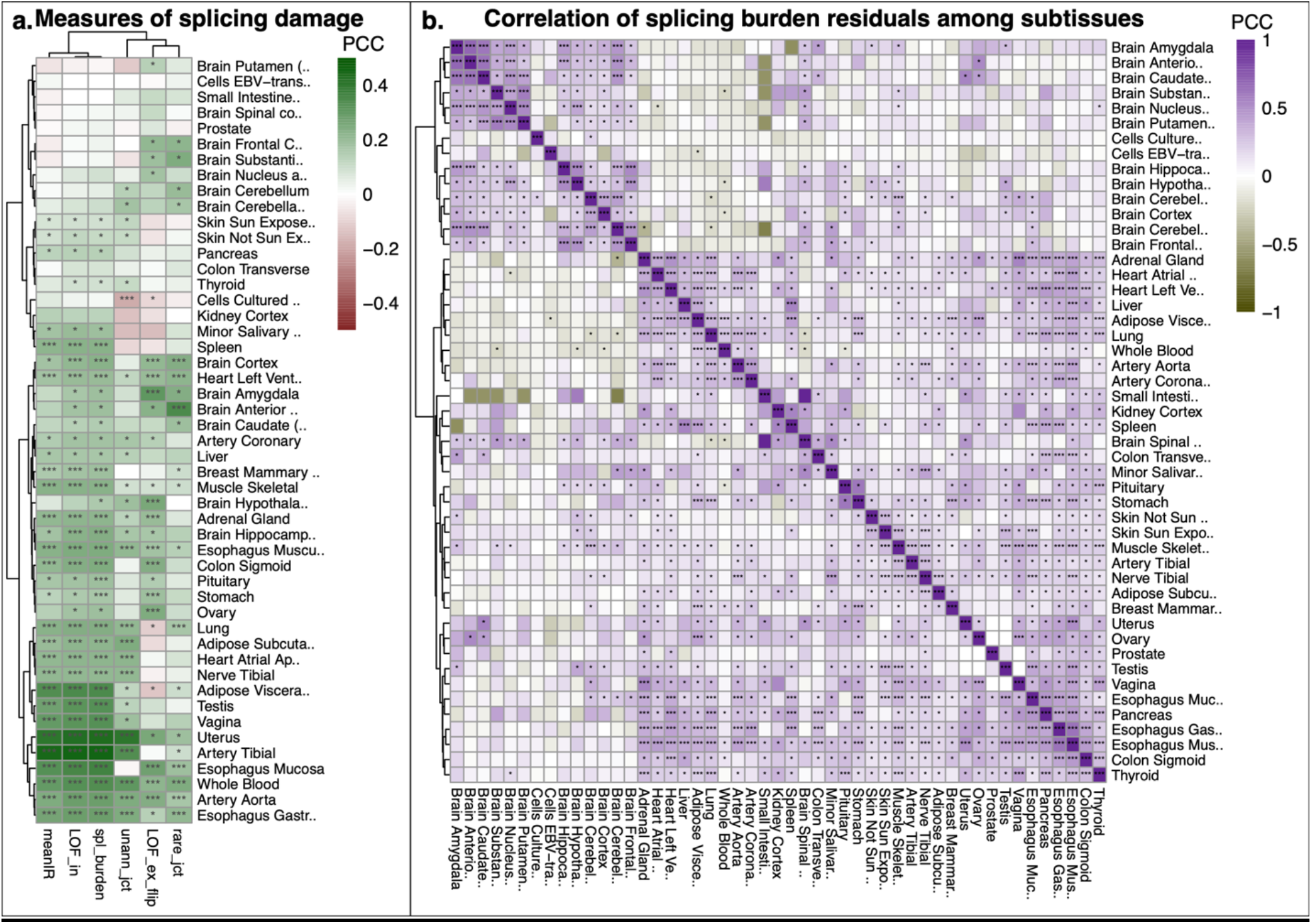
Measures of splicing damage. (a) Comparison of the various metrics introduced in this work, with colors representing their correlation with age in each subtissue (Pearson’s correlation coefficient). meanIR= mean PSI of IR events; LOF_in= mean PSI of LOF-inclusions (see Fig. 3a); LOF_ex_flip= mean (1-PSI) of LOF-exclusions; spl_burden= splicing burden; rare_jct= rare junction usage; unann_jct= unannotated junction usage. (b) Correlation of splicing burden residuals (after regression with age and cohort) across subjects, among all pairs of subtissues.

### Splicing damage correlates with age

We have shown that several measures of splicing noise increase with age in human tissues: rare junction usage, unannotated junction usage, and the mean IR level. We have also shown that, when analyzing IR, we should separate the events implicating an inclusion from those implicating an exclusion (relative to the functional mRNA isoform), due to the opposite interpretations of their PSI values, and opposite trend with age. We further defined one more measure of splicing noise, named “splicing burden”, as the mean of the observed frequencies of all IR events with LOF consequences (Fig. 3a), thus encompassing the PSI values of LOF-inclusions and the (1-PSI) values of LOF-deletions. This measure is highly correlated with mean IR, but it better captures the crucial difference of inclusions and exclusions. Among the aggregate measures presented in this work, it is the one that best correlates with age across human subtissues (Fig. 6a). Splicing burden represents the degree of impairment of functional mRNAs due to aberrant splicing, and thus it is well fit to reflect the concept of “splicing damage”.

A measure which reflects real biological damage should show correlation across tissues of the same individual. In fact, we expect that an individual with damage which is higher-than-average (for their age) in one organ would tend to have higher damage in other organs, too. This is because the exposure to many risk factors which are broadly detrimental (e.g. smoking, chronic diseases, genetic susceptibilities) is shared by the tissues of an affected individual. Therefore, we set to test whether we could detect such signal for our splicing burden measure. We calculated the residuals of the regression between splicing burden and age (including cohort as covariate), and then calculated the correlation of residuals among subtissues. In each pairwise comparison of subtissues, we considered only the subjects with available samples for both subtissues. Interestingly, we found that residuals were positively correlated among the majority of subtissues (Fig. 6b). This effect could not be explained by a correlation of residuals with age (<u>Fig. S12</u>).

### A novel molecular clock based on splicing damage can predict age

Epigenetic clocks are machine learning models to estimate the age based on DNA methylation levels ^25–29^. Notably, epigenetic clocks are said to capture the “biological age” rather than chronological age, meaning that exposure to various detrimental factors results in an observable acceleration of the predicted age. Despite their wide use, the nature of epigenetic clocks remains unclear.

We built a novel molecular clock explicitly based on measures of splicing damage. We prepared a feature table consisting of splicing burden components (i.e., PSI values of LOF-inclusions and LOF-exclusions). We then fit an elastic net regression model per subtissue, which allowed to select and appropriately weight those splicing events most informative for age. For 17 out of 49 subtissues, the resulting clock was “flat”: no feature was selected, so that the only model parameter was the intercept. We dismissed these subtissues, which were the ones with the weakest correlation between splicing burden and age. The rest of clocks are shown in Fig. 7a. As expected, age predictions on the test sets had greater correlation with age than splicing burden (i.e., the clock worked better than the simple mean of input features). Prediction error of splicing damage clocks was relatively high (∼10 years on average, Fig. 7a). Predicted ages spanned a narrow range, with larger discrepancies found in the youngest and the oldest samples (<u>Fig. S13</u>a). As in our previous analysis of splicing burden, we tested whether the residual signal (*deltaAge= predictedAge – realAge*) correlated among subtissues. In this case, delta age correlated negatively with age (<u>Fig. S13</u>b), which is not unusual for elastic net clocks. Since this would result in a deceiving correlation among subtissues, we considered the “age acceleration” instead ^29^, defined as the residual after fitting a *predictedAge ∼ age* linear model per subtissue (<u>Fig. S13</u>c). Remarkably, our analysis showed that, according to our splicing damage clocks, age acceleration positively correlates among the majority of subtissues (Fig.7 b-c).

**Fig. 7.**
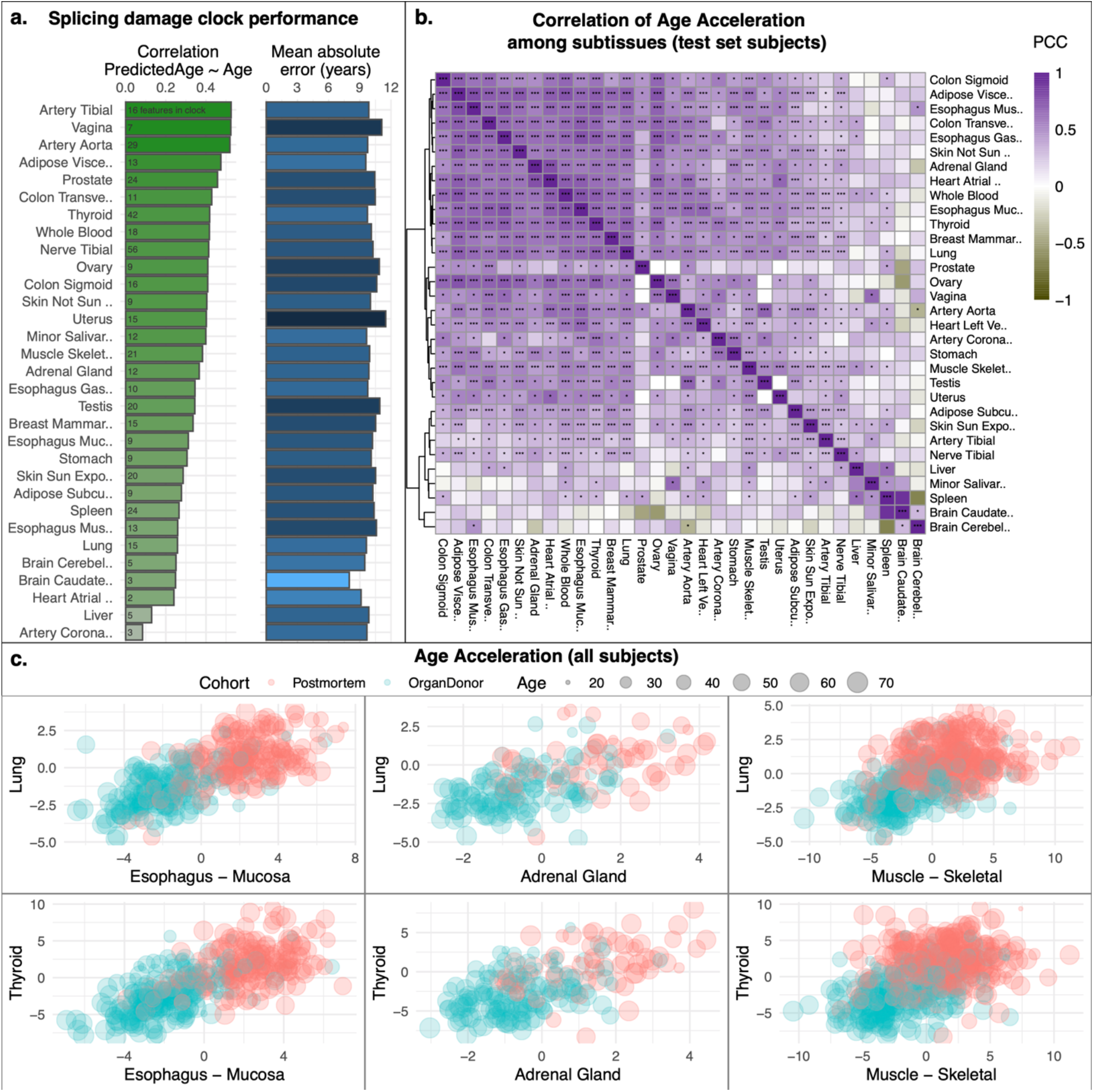
Molecular aging clock based on splicing damage. (a) Performance of our subtissue-specific clocks built with elastic net based on splicing damage features. It contains the Pearson correlation coefficient (green) and mean absolute error (blue) calculated on the test set, as well the number of features per clock (writing over green). See <u>Fig. S13</u> for scatterplots of raw data. (b) Correlation of age acceleration among subtissues of the same subjects, considering the test sets only. Stars within each square indicate statistical significance (***= FDR < 0.05; *= nominal p-value < 0.05). (c) Age acceleration scatterplots of some representative pairs of subtissues, where each point is a subject. The two cohorts feature distinct distributions of age acceleration values. Still, note that residuals are correlated across subtissues even considering each cohort separately.

### Splicing damage correlates with disease status

We further explored the relationship between splicing damage and disease. We manually selected 51 medical attributes recorded in GTEx as disease indicators (<u>Fig. S14</u>). These ranged from relatively mild features (e.g., influenza; unexplained cough) to serious illnesses (e.g., cancer; renal failure). The great majority of medical conditions were rare in the dataset, precluding analysis targeted to a single attribute. Instead, we calculated the total number of medical conditions per subject, collectively constituting a rough marker of disease status. This measure is likely subject to a high error, due to heterogeneity of diagnosis procedures among subjects, including lack of diagnosis for certain conditions. Still, we observed that it showed a strong and robust increase with age, as expected (Fig. 8a). When we correlated the number of conditions with splicing burden, we obtained a positive correlation for the majority of subtissues (Fig. 8c; positive correlation in 46 out of 49 subtissues; nominal p-value < 0.05 in 37; FDR < 0.05 in 24). Remarkably, this was not simply due to a common dependence on age: the residual of splicing burden (after regressing out age) also positively correlated with the number of conditions in most subtissues (Fig. 8b-c; positive correlation in 44 out of 49 subtissues; nominal p-value < 0.05 in 22; FDR < 0.05 in 9). This indicates that splicing damage is higher in the case of diseases, beyond what is explained by age alone.

**Fig. 8.**
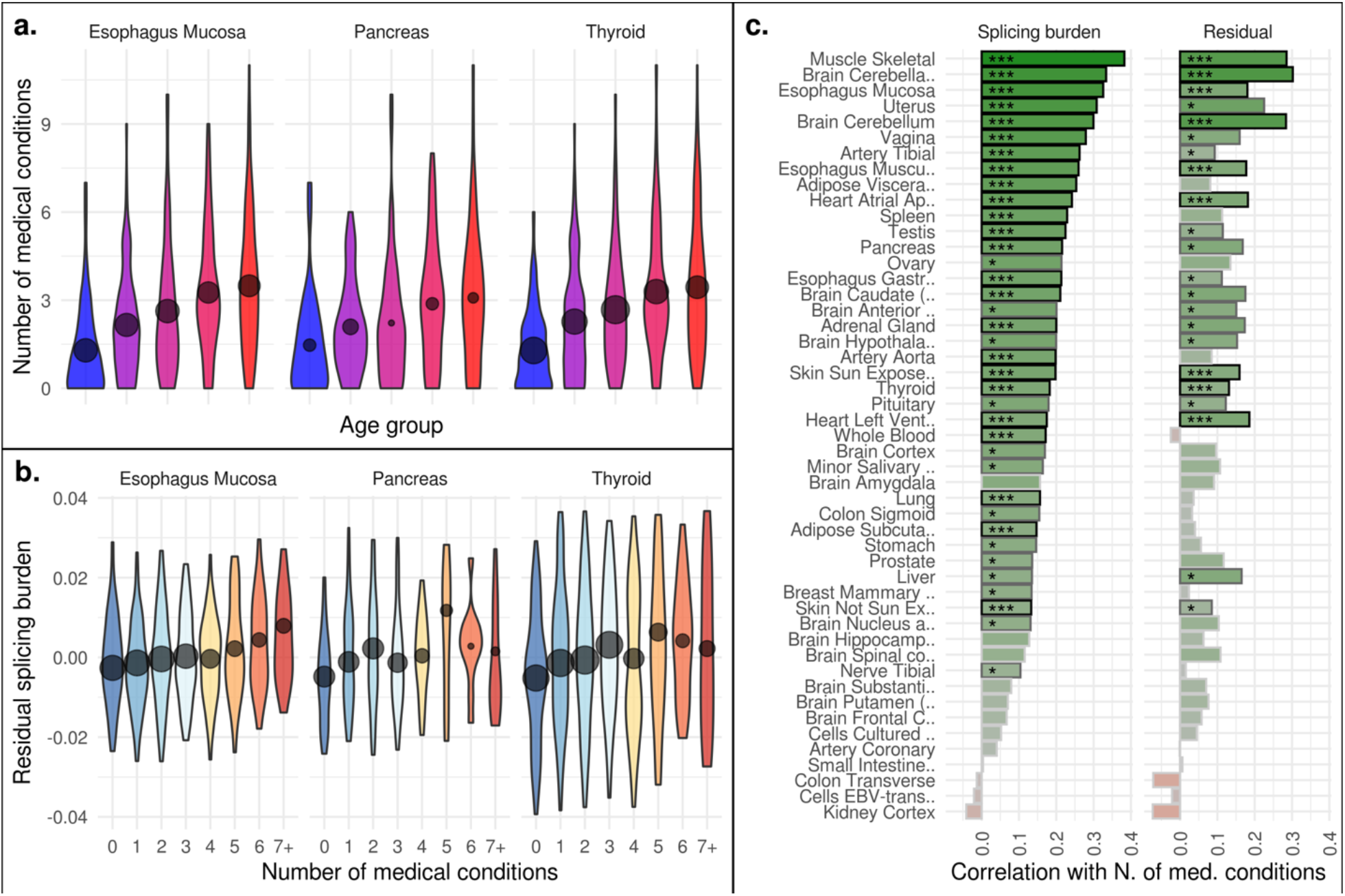
Splicing burden and medical conditions. (a) Distribution of the number of medical conditions per subject across age groups. This is shown for the subjects with available samples for three representative subtissues. (b) Distribution of residual splicing burden (after regressing out age) split by the number of medical conditions. (c) Pearson correlation coefficient between the number of medical conditions and splicing burden (first column) or its residual (second column). Stars indicate statistical significance (***= FDR < 0.05; *= nominal p-value < 0.05).

## Discussion

In this work, we analyzed age-related changes in splicing patterns across human tissues and found that the mean level of intron retention (IR) increases with age. Previously, IR increases with age were observed in human and mouse brain ^11,15^, fruit flies and *C*.*elegans* ^15–17^ However, by examining a remarkable diversity of human samples, we demonstrated that the age-related IR increase occurs consistently in most tissues. To our surprise, brain was among the tissues with the smallest effect. We also showed that IR correlates with overall disease status, even beyond its dependence on age. These observations indicate that the IR increase during aging is not a specific feature of the brain or neurodegeneration, but instead is a more general indicator of disease.

Much of our work involved the analysis of IR events quantified by RNAseq. An important feature in this type of data (to our knowledge, unaccounted for in previous research) is that it encompasses events with opposite functional interpretations (Fig. 3a-b): inclusions (IR-in) and exclusions (IR-ex). Inclusions constituted the great majority of IR events, and corresponded to the common intuition of IR (Fig. 1a). These tend to increase with age in most tissues, leading to a progressively higher representation of dysfunctional mRNAs with impaired protein products (<u>Fig. S2</u>). This indicates that “genuine” introns fail to be recognized or processed, which we interpret as a sign of declining splicing efficiency with age.

Exclusions, on the other hand, correspond to the removal of functional mRNA regions. This also leads to transcripts with impaired coding sequences, but by subtraction rather than addition. Strikingly, exclusions also show higher incidence with age (i.e., decreased PSI) (Fig. 3b, <u>Fig. S4</u>b). We interpret this as a sign of decline in splicing specificity: more and more spurious regions are spliced out with age. To confirm this, we applied a second approach to separately analyze exon-exon junctions *de novo*. We found that the usage of rare and unannotated junctions increases with age in most human tissues (Fig. 6a, <u>Fig. S11</u>), supporting our hypothesis. The decline in both splicing efficiency and specificity leads to the general deterioration of the transcriptome, as functional mRNAs are increasingly diluted with dysfunctional isoforms. This represents the accumulation of splicing damage, and to measure it, we defined a “splicing burden” index, which we found to positively correlate with age in almost all tissues (Fig. 6a).

Our results are consistent with the reports of altered splicing with age ^9,11,15,17^, but pinpoint two concurrent modalities of mis-regulation. Additionally, we assessed tissue heterogeneity as a potential explanatory factor. When a tissue is sampled by bulk RNAseq, an apparent pattern of altered splicing could be due to the infiltration by other cell types, in which distinct transcript isoforms are prominent. However, we observed that the age trend of individual IR events is quite consistent across tissues. This undermines tissue heterogeneity as an explanation for the age-related increase of IR, and points instead to molecular processes occurring within cells.

We also observed some differences among tissues. Most notably, mean IR correlates with age more poorly in brain than other tissues. We wondered whether this could be attributed to the different age distributions, e.g., since all brain samples belonged to a cohort (postmortem) notably older than the other (organ donor) (<u>Fig. S1</u>). However, when we analyzed splicing burden in postmortem samples only (with nearly homogenous age distributions across subtissues, <u>Fig. S1</u>), we obtained essentially the same tissue-specific pattern (<u>Fig. S15</u>). This suggests that differences among subtissues may be of biological nature. Interestingly, cultured cells derived from GTEx samples (fibroblasts and EBV-transformed lymphocytes) appeared as outliers from the actual tissues. In cell cultures, the correlation between splicing burden and age is poor (Fig. 6a), the burden residual does not correlate across subtissues (Fig. 6b), and individual IR events are mostly unchanged with age (Fig. 2). This suggests that the age-dependent signal of splicing damage is lost in cell culture, so these systems may not be good models to study this phenomenon.

The mechanism of splicing deterioration with age is yet unknown, but a link with splicing machinery dysregulation was implied. This is based on the observation that several genes in this pathway showed altered expression with age, both in absolute levels and in isoform usage ^6,8,9,15,30^. In our study, rather than seeking pathways overrepresented with the genes with altered splicing, we identified pathways whose expression correlated with global indicators of splicing damage, such the mean IR per sample. Our regression analysis (Fig. 5) revealed processes previously implicated in aging (e.g., DNA repair, translation, protein degradation), despite the fact that we controlled for age in our model design. Notably, a considerable proportion of spliceosome factors correlates negatively with IR. This reinforces the possibility of a direct link between splicing machinery levels and declining splicing fidelity with age. Within the spliceosome, the majority of the Prp19 complex had a significant negative correlation with IR. This complex constitutes a major player all throughout the splicing process, and was also involved in genome maintenance, protein degradation, biogenesis of lipid droplets, transcription elongation and mRNA export ^31^. Since it engages in multiple molecular processes involved in age-dependent decline, we propose the Prp19 complex as a potentially important actor in aging, or even a possible target for anti-aging treatments. Notably, a recent report on GenAge ^32^ shows that the overexpression of the Prp19 ortholog significantly extended lifespan of female fruit flies, by up to 25% ^33^.

We were surprised to discover roughly the same pathways correlated with inclusions and exclusions (Fig. 5, <u>Fig. S10</u>). It is possible that the decline in splicing efficiency and specificity may be the two sides of the same coin. An attractive possibility is that, whether we measure one or the other (or any form of splicing damage), we capture a general index of molecular dysfunction, i.e. it reflects biological age. There are three additional observations supporting this notion. First, the residual of splicing damage with age correlates among most tissues, whether we quantify it as splicing burden (Fig. 6) or through the elastic net clock (Fig. 7). The residual represents the surplus (or deficit) of damage of a given individual compared to the population mean of the same age. In general, we think of damage as increasing with age, but of course there are many factors (e.g., lifestyle, injuries, treatments) which can accelerate this process or slow it down, creating a gap between chronological age and biological age. We expect these factors to exhibit generally concordant effects across tissues. Thus, a correlation of residuals is precisely what we expect from a measure capturing biological damage, and it was remarkable to observe it for our splicing-based metrics (though we must note that such correlation may have other contributions, e.g., due to sample structure). Second, we observed that IR-correlated pathways largely overlapped with those changing with age, despite the fact that we corrected for age (Fig. 5, <u>Fig. S9</u>). This may suggest that individuals with higher splicing damage are “biologically older”, which is reflected in their global expression profile --roughly along the same axis of transcriptomic change taken by natural aging. Third, we observed that, in most tissues, splicing burden positively correlates with disease status (number of medical conditions), even after correcting for age (Fig. 8). All these observations support splicing damage measures as biologically relevant indicators of deterioration in molecular function.

Although age can be estimated from splicing damage features, the age predictions were not as accurate as human clocks based on DNA methylation ^25,34–38^ or other molecular features ^39–42^. This is not entirely surprising, as other clocks also capture processes other than aging, most notably neutral changes and changes associated with adaptation to damage. On the other hand, splicing damage has the advantage of explicitly measuring a form of molecular damage (transcriptome deterioration due to mis-splicing), so that it directly reflects the nature of aging as a progressive accumulation of damage. We believe that approaching aging in explicit terms of observable molecular damage is compelling to better understand this process, quantify its effects, and ultimately find methods to attenuate it or reverse it. Accordingly, this could also lead to the development of better clocks that capture the nature of aging, as opposed to age-related changes, age-related diseases or age-related mortality.

A relevant open question is whether splicing dysregulation is itself causative of some aging phenotypes ^6^. Interestingly, there are indications from studies in *C*.*elegans* that it may be the case. Splicing factors SFA-1 and HRPU-1 have been shown to mediate the effect of dietary restriction on lifespan extension ^16,17^. Strikingly, SFA-1 overexpression is sufficient to significantly extend *C*.*elegans* lifespan ^17^. Further functional studies will be instrumental to pinpoint the precise role of splicing machinery in aging, and perhaps identify actionable targets to extend the lifespan and healthspan of vertebrates, too.

An age-related increase in molecular damage is the cornerstone of aging ^1^. This damage manifests in many forms, such as damage to metabolites, DNA and proteins ^43^. Although gene expression analyses have been a major tool in examining the underlying biology of aging and available data far exceeds other “omics” (e.g. proteome or metabolome profiling), little information has been reported on the age-related damage to the transcriptome. In part, this is due to the difficulty of assessing chemical modifications in the mRNA through sequencing or microarray analyses. In this regard, our current study resolves this major roadblock by showing that the transcriptome damage may be quantified from RNAseq data in the form of intron retention and spurious splicing. Importantly, this damage represents transcriptome deterioration, as opposed to age-related changes in gene expression as in the traditional RNAseq analyses, which include, to a large degree, responses to age-related damage.

## Methods

### RNAseq data and gene expression

We analyzed RNAseq samples in GTEx v8 ^18^, filtering out the subtissues with fewer than 50 samples. The resulting set consisted of 17,329 samples encompassing 49 subtissues of 948 subjects (Fig. 1a). Expression profiles at gene and transcript level in Reads Per Kilobase of transcript, per Million mapped reads (RPKM) and transcripts-per-million (TPM), respectively, were downloaded from the GTEx portal. These are based on the Gencode v26 annotation and the GRCh38.p10 genome assembly. For each subtissue, samples were split in 5 age groups of approximately equal size (quintiles) using R function quantcut.

### Splicing quantification

We used two different methods for splicing quantification. The first is the program Suppa v2 ^19^, which deconvolutes transcript quantifications to infer PSI (percent-spliced-in) values representing the frequency of every splicing event derived from a genomic annotation (Gencode v26). We used Suppa to quantify all “retained intron” events, wherein a PSI value of 0 signifies complete splicing, and 1 signifies complete retention (IR). We filtered out events with an apparent intron length shorter than 10bp. In Suppa, any IR event for which all transcripts defining it do not pass an expression level threshold (<1 TPM) are not quantified, and are assigned “NA” values. We thus filtered IR events to focus on those quantified in sufficient samples for meaningful analysis. For each subtissue, we generated sets of IR events by applying a “coverage filter” of 80%, 90% and 100%, so that an event was included only if quantified in at least that proportion of samples in each age group (to avoid potential age-related bias). The number of IR events in these sets is shown in Fig. 1b.

We also used a second method for splicing quantification: Leafcutter, which employs split-mapped reads to quantify exon junctions *de novo*. In contrast to Suppa, this program can detect unannotated events, but it cannot be used to quantify IR. Leafcutter quantifications (absolute read counts) were downloaded from the GTEx portal and used as described in the text.

### Classification of IR events by their functional consequence

We selected a reference mRNA isoform per gene, and then inferred the alteration of its protein product upon occurrence of each IR event in our dataset. Annotations of “principal” transcripts were obtained from the APPRIS database ^20^, release version 2017_06.v23.HS.e88v22. For genes with more than one APPRIS principal isoform, we selected the one with the highest mean expression level in the subtissue under analysis. We defined 11 possible types of functional consequences. The most abundant type consisted in a mRNA “inclusion” (i.e., the retained intron) resulting in the introduction of a frameshift (“in-FS”). The second-most abundant type were inclusions maintaining the frame, but introducing a premature stop codon (“in-PTC”). In few cases, inclusions maintained the frame without additional stops, thereby adding a peptide to the protein sequence (“in-pep”). In other cases, the retained intron occurred within the 5’ or 3’ UTR (“in-UTR”). In all other cases, the coordinates of the IR event did not correspond to any of the introns of the reference transcript. From these, we isolated those events mapping entirely within a reference exon: these events consist in removal of portions of the functional transcript upon splicing, and we referred to them as “exclusions”. Most exclusion events introduced frameshifts (“ex-FS”). Others removed the region corresponding to the start (“ex-Nter”) or the end (“ex-Cter”) of the coding sequence. Others eliminated coding regions but maintained the frame (“ex-pep”) or mapped entirely in the 5’ or 3’ UTR (“ex-UTR”). The category “other” comprises all IR events not qualifying for any of the previous definitions; for these, none of the two alternative outcomes (splicing / retention) results in production of the reference mRNA, so that they are of unclear interpretation. Lastly, the category “noncoding” was used for all IR events in genes with no principal isoforms annotated, since these corresponded to non-coding genes.

These consequence types were further grouped in categories, as depicted in Fig. 3a. Besides, a separate set of constitutive introns was used to analyze the sequence determinants of IR age trends. Constitutive introns were defined as all those present in reference transcripts (as defined above), excluding those annotated as retained in any isoform (i.e., those introns quantified by Suppa without any coverage cut-off).

### Linear models, gene expression regression, and gene set enrichment analysis

Linear models were built in R with lm function from base package stats, and linear mixed models were built using lmer from the packages lme4 and lmerTest. Gene expression regression analysis instead was performed with Voom-Limma ^44,45^ and edgeR ^46^. Gene set enrichment analysis was performed using Clusterprofiler ^47^, ranking genes by the t-value assigned by Limma. We limited our analysis to KEGG pathways. To obtain the list of pathways with significant and consistent effects across subtissues, we filtered those with FDR <0.05 and the same direction (positive or negative) in more than half of subtissues (see <u>Fig. S7</u>).

### Splicing damage clock

We prepared a feature table with the PSI values of individual LOF-inclusion and LOF-deletion events (Fig. 3a), quantified per sample. We randomly selected 30% of the 948 subjects as potential test subjects. We partitioned samples to train and test sets for each subtissue, separately, following the split at the level of subjects. Then, we trained and tested an elastic net regression model for each subtissue. The lambda parameter was optimized on the given training set by the built-in 10-fold cross-validation of the Python package Glmnet (https://github.com/civisanalytics/python-glmnet, v2.2.1; alpha=0.5, n_splits=10). We provide the intercepts and the non-zero weights of the clocks as well as the predicted ages (i.e. ‘splicing ages’) of the subject in Supplementary Data D1.

### Clustering, statistical analysis, and plotting

All data analysis and statistical tests were performed in R. Multiple test corrections were performed by the Benjamini-Hochberg procedure (i.e., FDR= false discovery rate). Hierarchical clustering was performed using Euclidean distance matrices using the “Ward.D2” method. Clustered heatmaps were produced with the pheatmap package; all other plots were produced using the ggplot2 package ^48^.

## Supporting information

All supplementary figures

## Acknowledgements

Supported by NIH grants to VNG. MM was supported by grants PID2020-115122GA-I00 and RYC2019-027746-I by the Spanish Ministry of Science and Innovation.

