## Supplementary material for "Deterioration of the human transcriptome with age due to increasing intron retention and spurious splicing": All supplementary figures

### Supplementary data

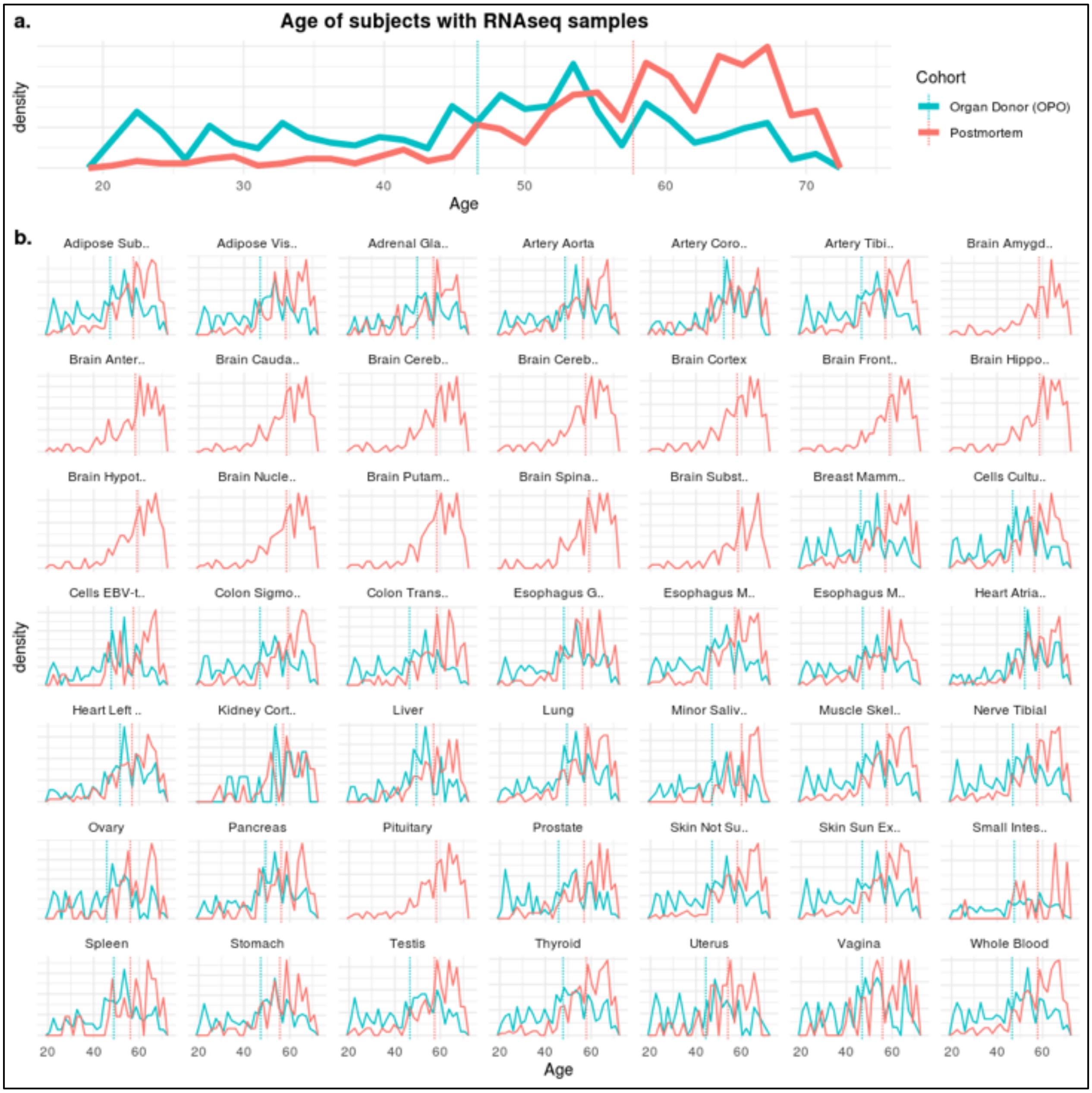


**Fig. S1**. Age profile of GTEx cohorts. Panel (a) shows the profile for all unique subjects with at least one RNAseq sample, while panel (b) shows this information per subtissue. Note that the cohort "Surgical" is omitted here since it represents less than 0.5% of samples analyzed (see Fig. 1).

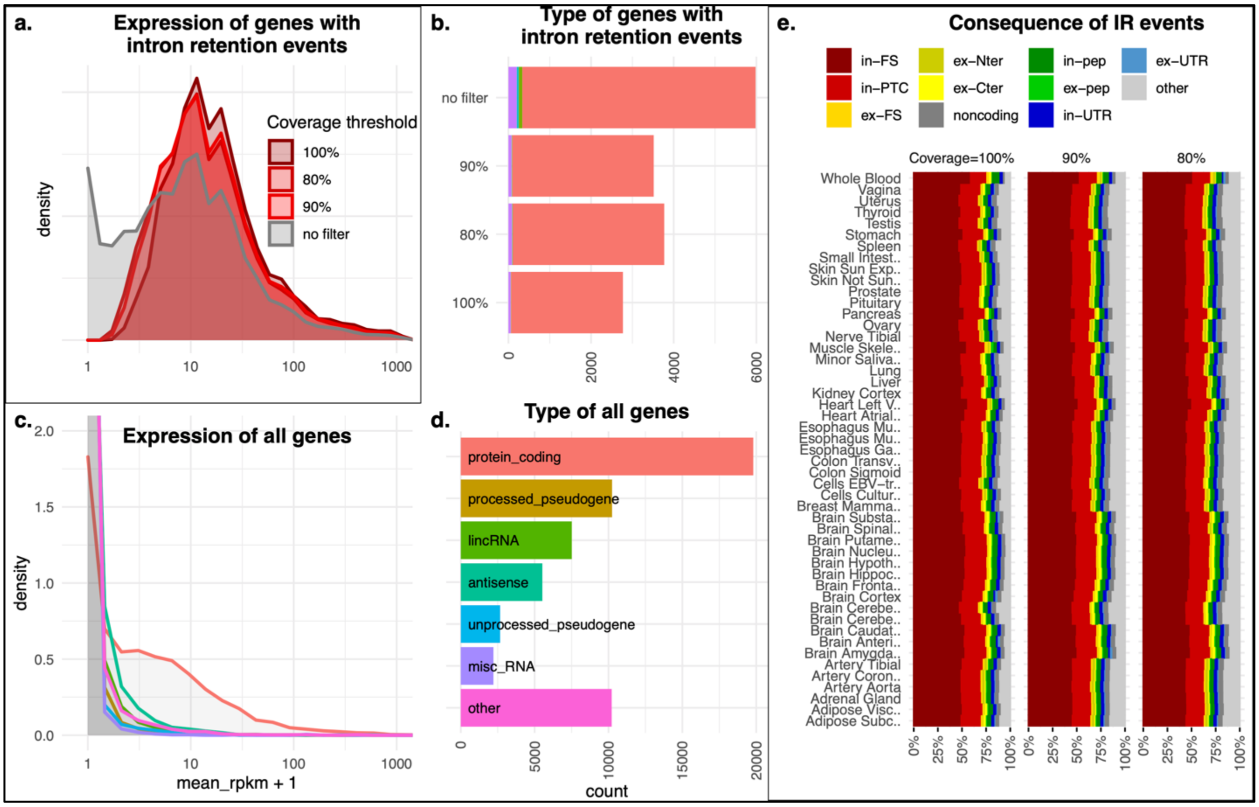


**<u>Fig. S2</u>**. Effect of coverage filter on the expression and type of genes with intron retention events. (a) Distribution of expression values of genes with IR events passing the coverage threshold. (c) The same for all genes. (b) and (d) show the types of genes, with or without coverage filtering. Panels (b), (c) and (d) have the same color coding, shown in panel (d). (e) Functional consequences assigned to IR events (see <u>Fig. 3a</u>).

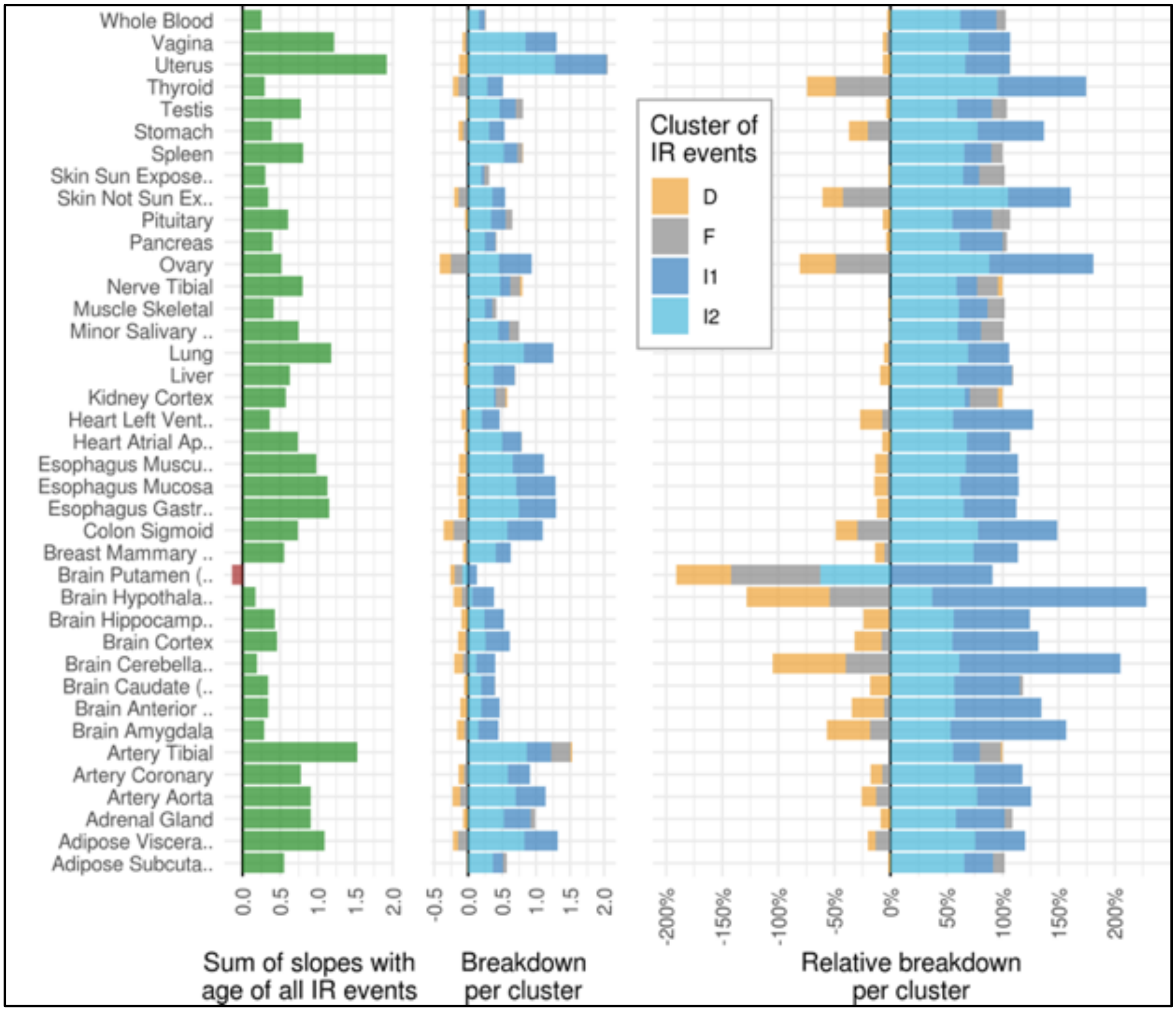


**Fig. S3**. Breakdown of slopes with age per IR event cluster. We computed the sum of PSI ~ age slopes for all events (first column), or separately per cluster (second column). The third column shows the relative values per cluster normalized by the total of all events. This value retains the sign (so in the presence of a cluster with a negative contribution, the total of breakdowns in absolute value exceeds 100% for that subtissue). For visualization purposes, the subtissues with the total sum of slope smaller than 0.12 in absolute value were excluded.

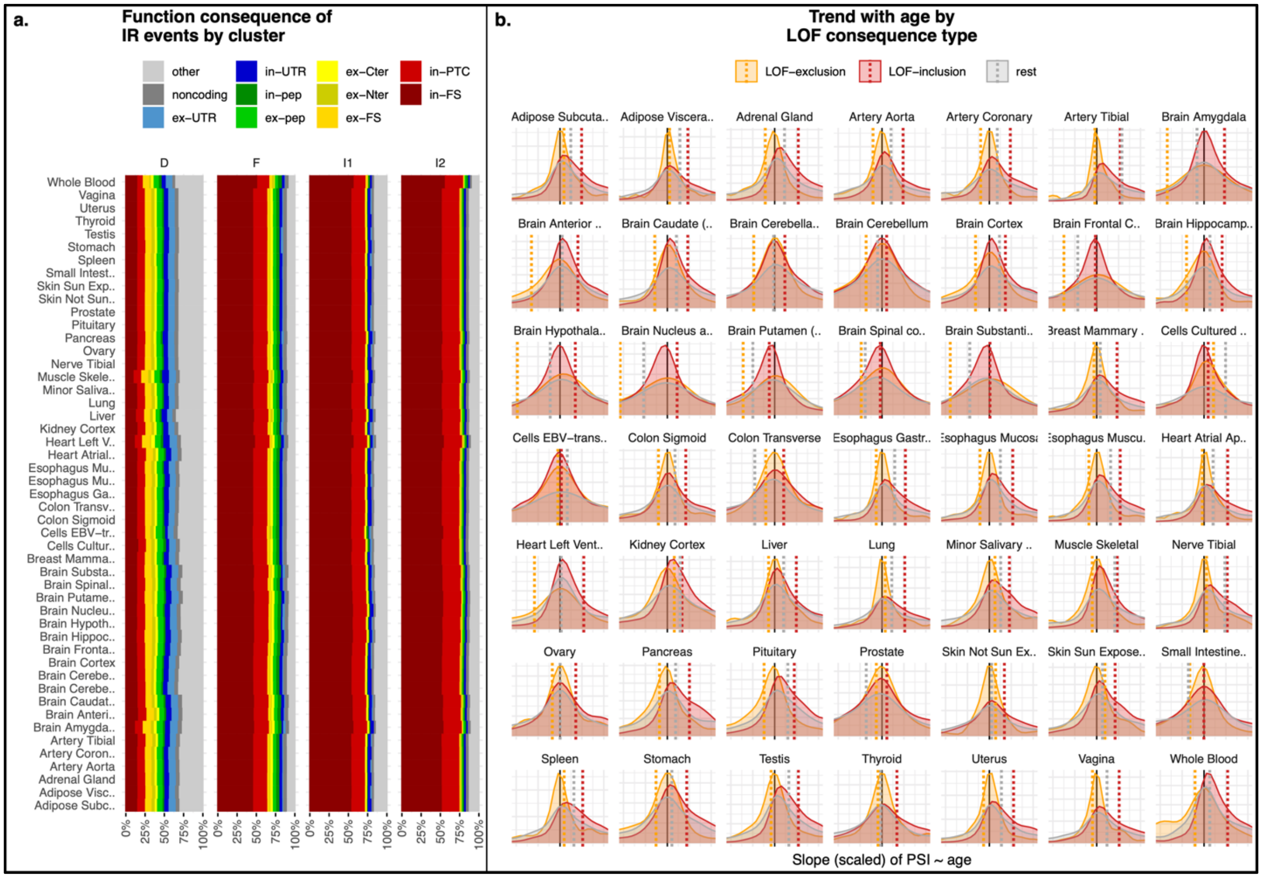


**<u>Fig. S4</u>**. Age changes of IR events with different functional consequences. (a) Composition of different clusters of IR events in terms of their functional consequences. IR events were assigned to the clusters designed as D (decrease), F (flat), I1 (strong increase) or I2 (increase) based on the slope of their PSI value with age (see Fig. 2a). Functional consequences per event (colors) were assigned based on their effect on the gene structure of a reference transcript per gene (see Fig. 3a). (b) Age distribution of slope of LOF events. We fit a linear model PSI ~ age for each IR event, and considered the resulting slope. The plot shows the density of slopes separately for the LOF-inclusions events (including "in-FS" and "in-PTC"), the LOF-exclusions events (including "ex-Nter", "ex-Cter", and "ex-FS") and the "rest" of events. Dotted vertical lines show the mean slope per category, and the vertical black line marks zero. For visualization purposes, the X-axis range was scaled per subtissue, symmetrically around zero, to the 80% percentile value.

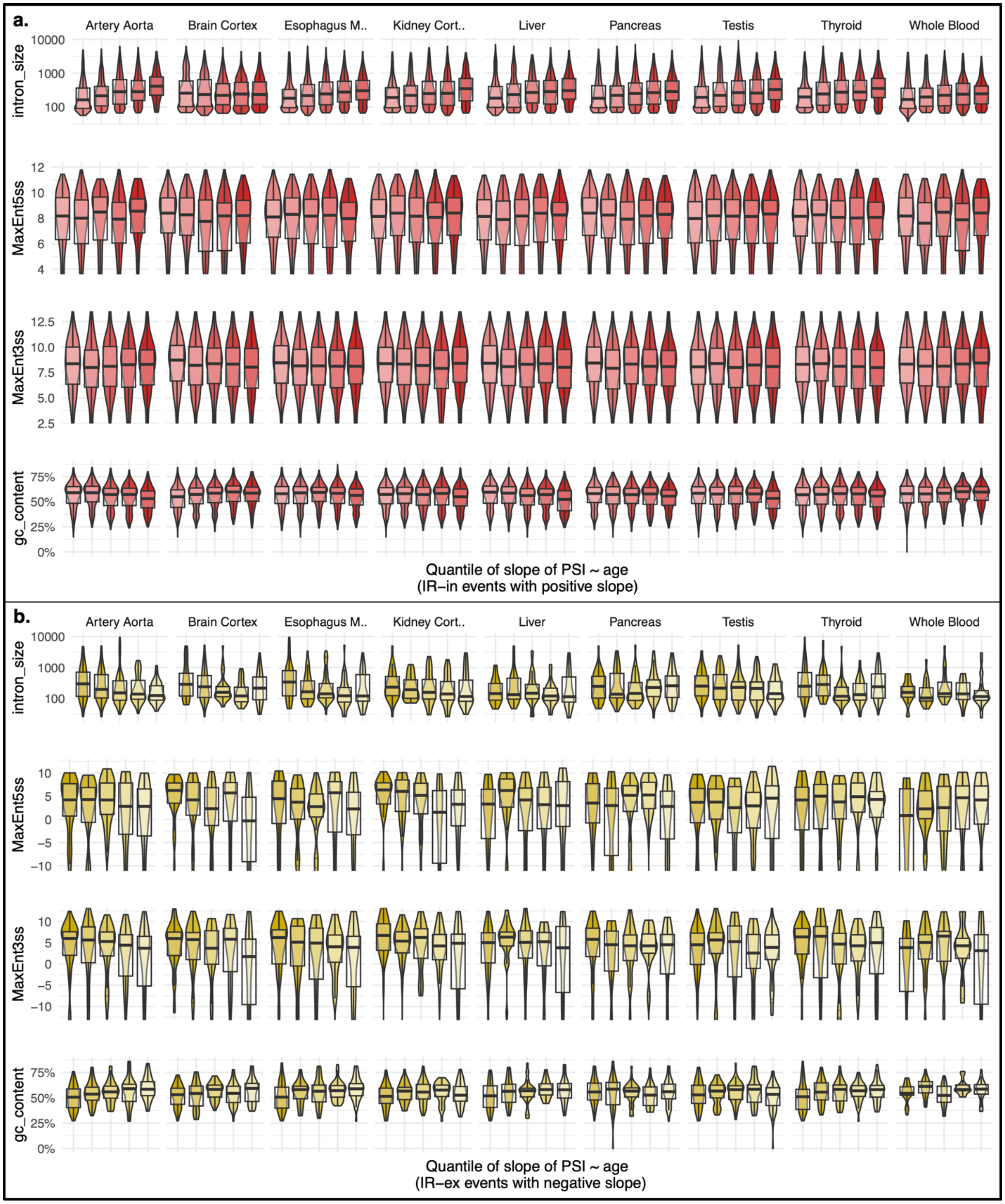


**Fig. S5**. Determinants of intron retention change with age. The figure shows the distribution of intron features (intron size, splice site scores for 5’ and 3’, and GC content) in quantiles of PSI ~ age slopes. (a) Age-increasing IR-in events, wherein the fifth quantile corresponds to introns with the largest increase. (b) Age-decreasing IR-ex events, wherein the first quantile corresponds to the largest decrease.

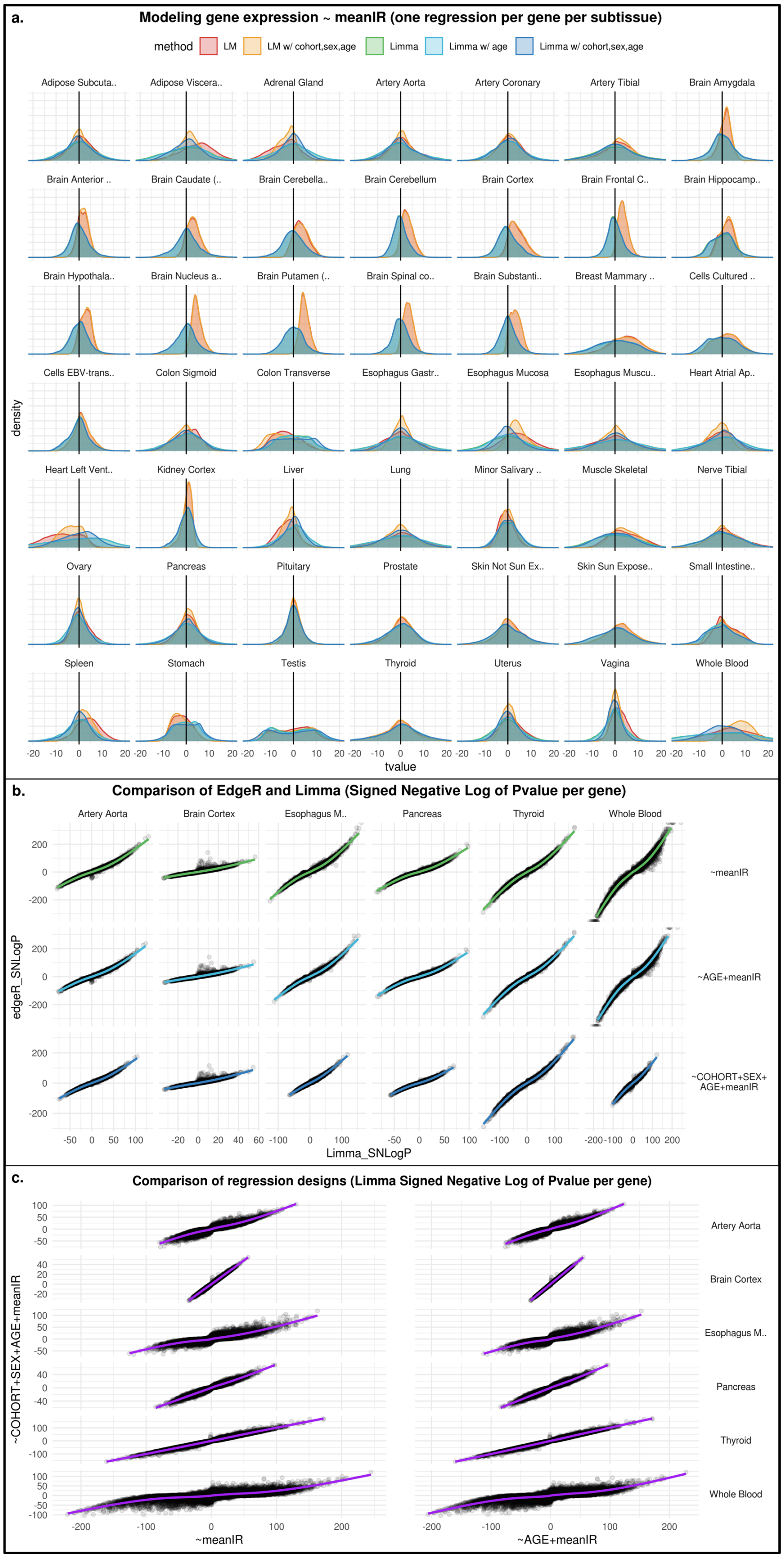


*(Fig. S6; Caption on next page)*

**Fig. S6** (previous page). Regression of gene expression and mean intron retention. Panel (a) shows the distribution of t-values associated with the test of the slope of meanIR / geneExpression being different than zero, comparing different methods (LM= simple linear model, Limma= Voom-Limma; see Methods) and various designs differing by the covariates considered (age, cohort, sex). Note that, in many subtissues, the distribution of t-values is skewed for simple linear models; taken at face value, this would imply that the majority of genes exhibit a significant correlation with meanIR, but it is rather an effect of a shift in the shape of gene expression distribution with age (see Fig. 4). Presumably due to the normalization employed (TMM), the Limma method rescued this issue (with only testis still showing an unusual t-value distribution). (b) Comparison of two algorithms to regress gene expression with a continuous variable (meanIR): Limma and edgeR. Each point represents a gene, with coordinates determined by their SNLogP (negative log of the p-value associated to the slope between gene expression and meanIR, with sign to indicate positive or negative correlation). While Limma is more conservative, the two algorithms are essentially in agreement. Analogously, panel (c) shows a per-gene comparison of Limma results across model designs differing by the co-variates included.

**Fig. S7** (next page). KEGG pathways enriched in genes whose expression correlates with mean IR, using cohort, sex and age as covariates. Rows represent pathways, columns represent subtissues, and both dimensions were subject to hierarchical clustering. Each square is colored by the normalized enrichment score (NES), and contains white text indicating statistical significance (***= FDR < 0.05; *= nominal p-value < 0.05). We selected those pathways showing a consistent effect across subtissues (FDR < 0.05 and same direction in >50% of subtissues), marked in purple on the left side of the plot. These pathways are further shown in Fig. 5.

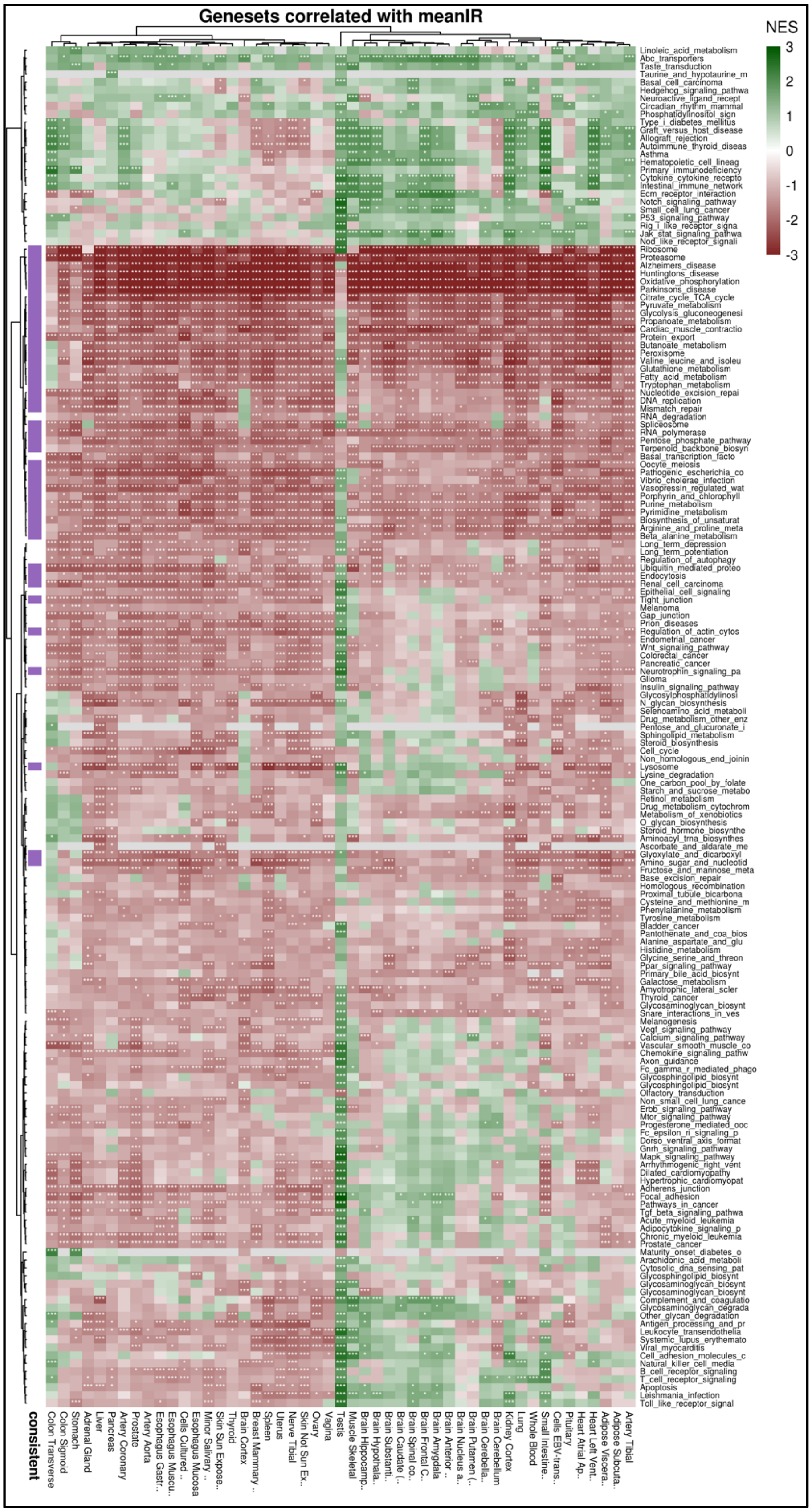


*(Fig. S7; Caption on previous page)*

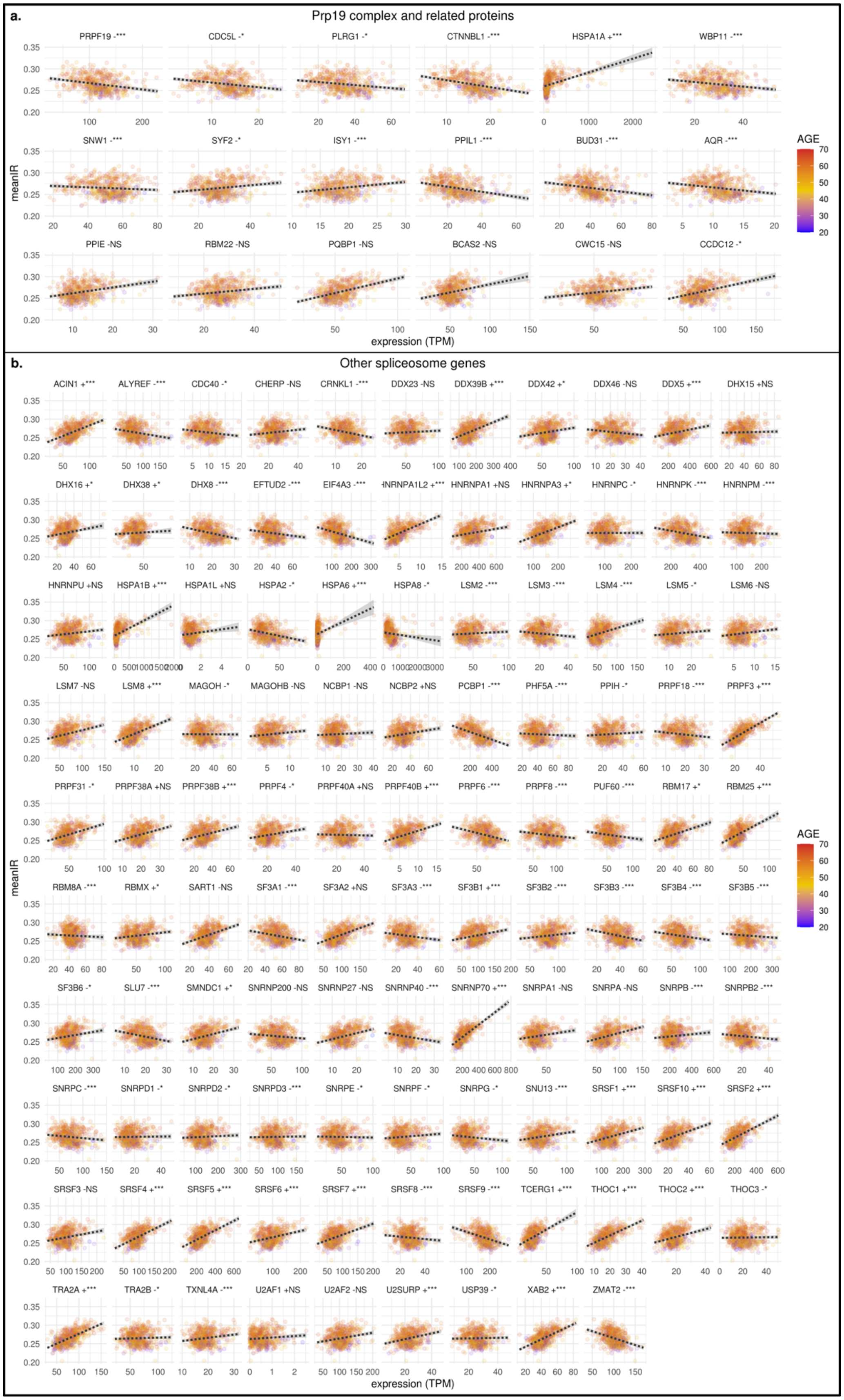


*(Fig. S8; Caption on next page)*

**<u>Fig. S8</u>** (previous page). Correlation between expression of spliceosome genes and mean intron retention per sample. (a) Genes annotated as a part of the Prp19 complex or related to it. (b) All other genes annotated in KEGG pathway for the spliceosome. Expression values were taken from the representative tissue of Esophagus mucosa. Next to each gene name, a “+” or “-” indicates the sign of the relationship, followed by a symbol to indicate statistical significance (NS for not significant, * for p-value <0.05, *** for FDR <0.05). Note that, since the relationship between these variables was determined in a model accounting for additional co-variates (cohort, sex, age), the sign shown does not always correspond to that implied by the trend line shown (which only depend on expression and meanIR).

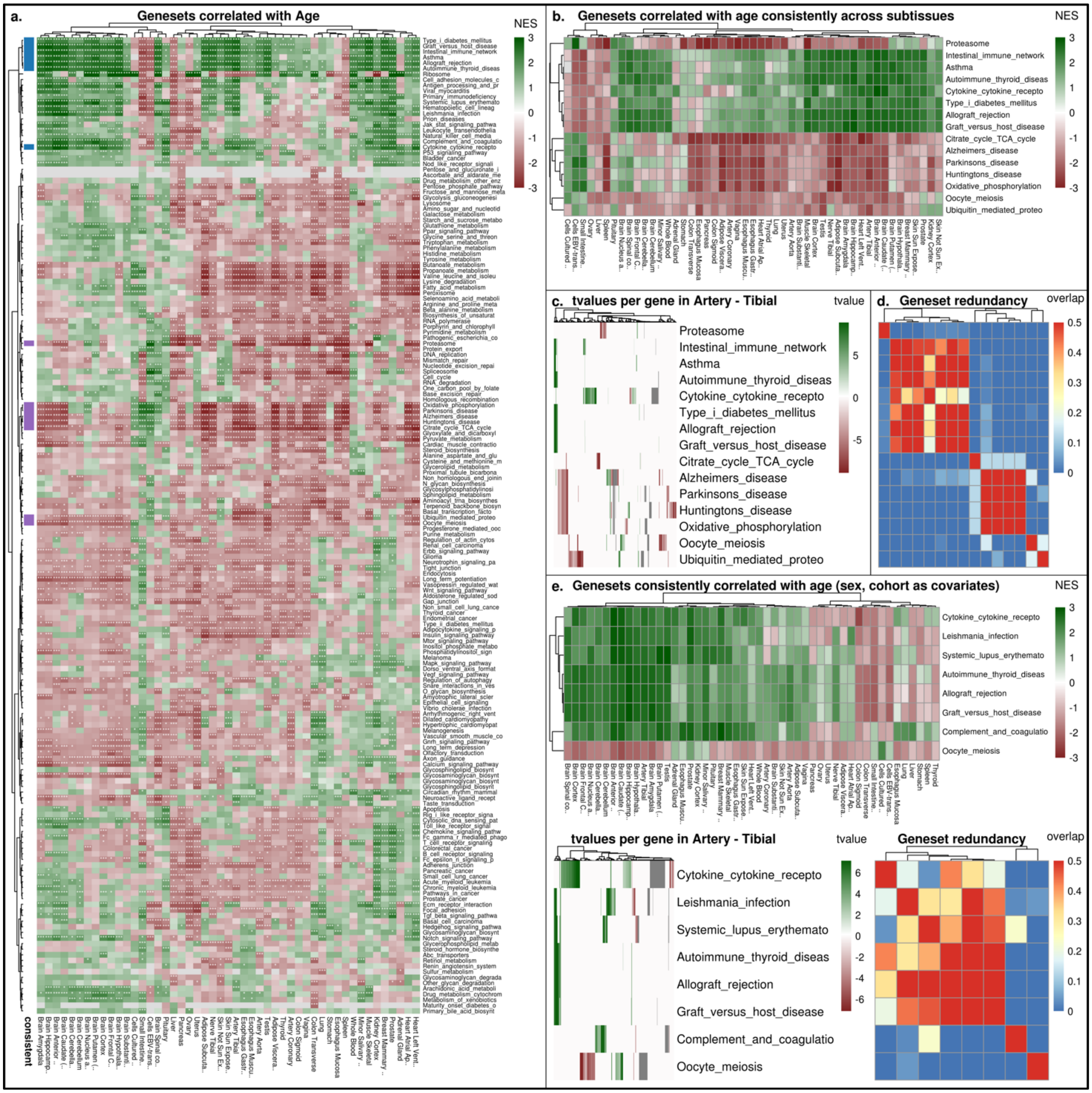


**Fig. S9**. Pathway enrichment analysis of genes whose expression correlates with age. The plots presented are analogous to Fig. 5, but here the model design used was 'expression ~ age' (panels a-d) or 'expression ~ cohort + sex + age' (panel e).

**Fig. S10** (next page). Pathway enrichment analysis of genes whose expression correlates with mean PSI of IR-in or IR-ex events. The figure shows the results of our analysis, analogous to that performed for mean IR shown in Fig. 5, but here considering instead the mean PSI value of IR-in events (panel a) or 1 - the mean PSI of IR-ex events (panel b). We flipped the PSI value of mean IR-ex (i.e., transformed as x'=1-x) to have the same conceptual direction, with greater values corresponding to increasing damage.

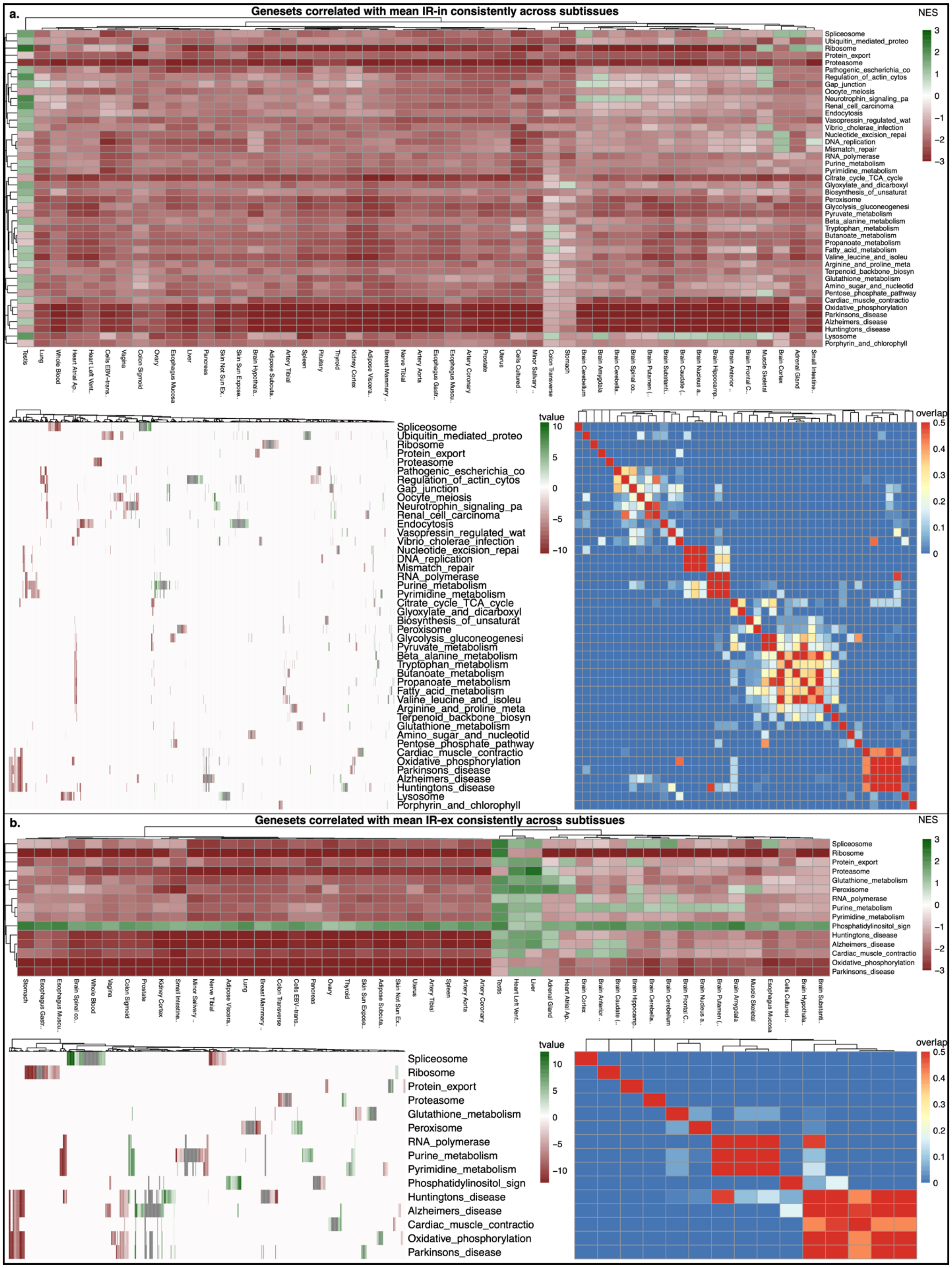

(Fig. S10; Caption on previous page)

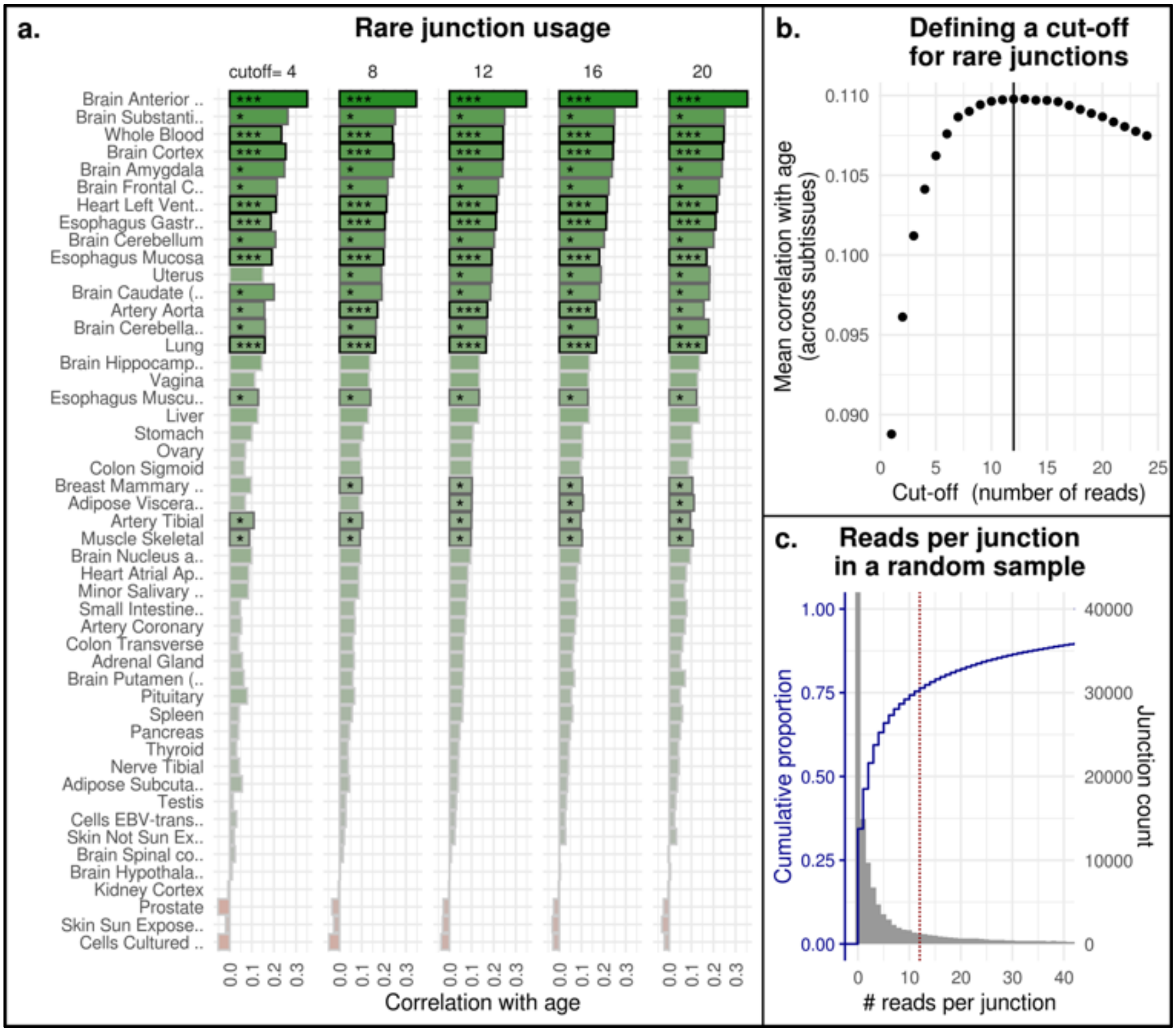


**Fig. S11**. Definition of rare junction usage and correlation with age. (a) Pearson correlation coefficient between rare junction usage and age. Rows represent different subtissues, and columns different cut-offs to define rare junction (i.e., a rare junction is one that is supported by not more than X Leafcutter reads). (b) The panel shows how the correlation of this metrics as a function of age (mean across subtissues) varies with the chosen cut-off. The correlation is robust across a wide range of the cut-off parameter. We chose the cut-off value of 12, which maximizes the mean correlation with age, for further analysis. (c) This panel shows how Leafcutter reads are distributed across junctions: the majority of reported junctions are supported by few reads. Yet, the total amount of reads in these rare junctions represents only a minor fraction of all reads (4-10%).

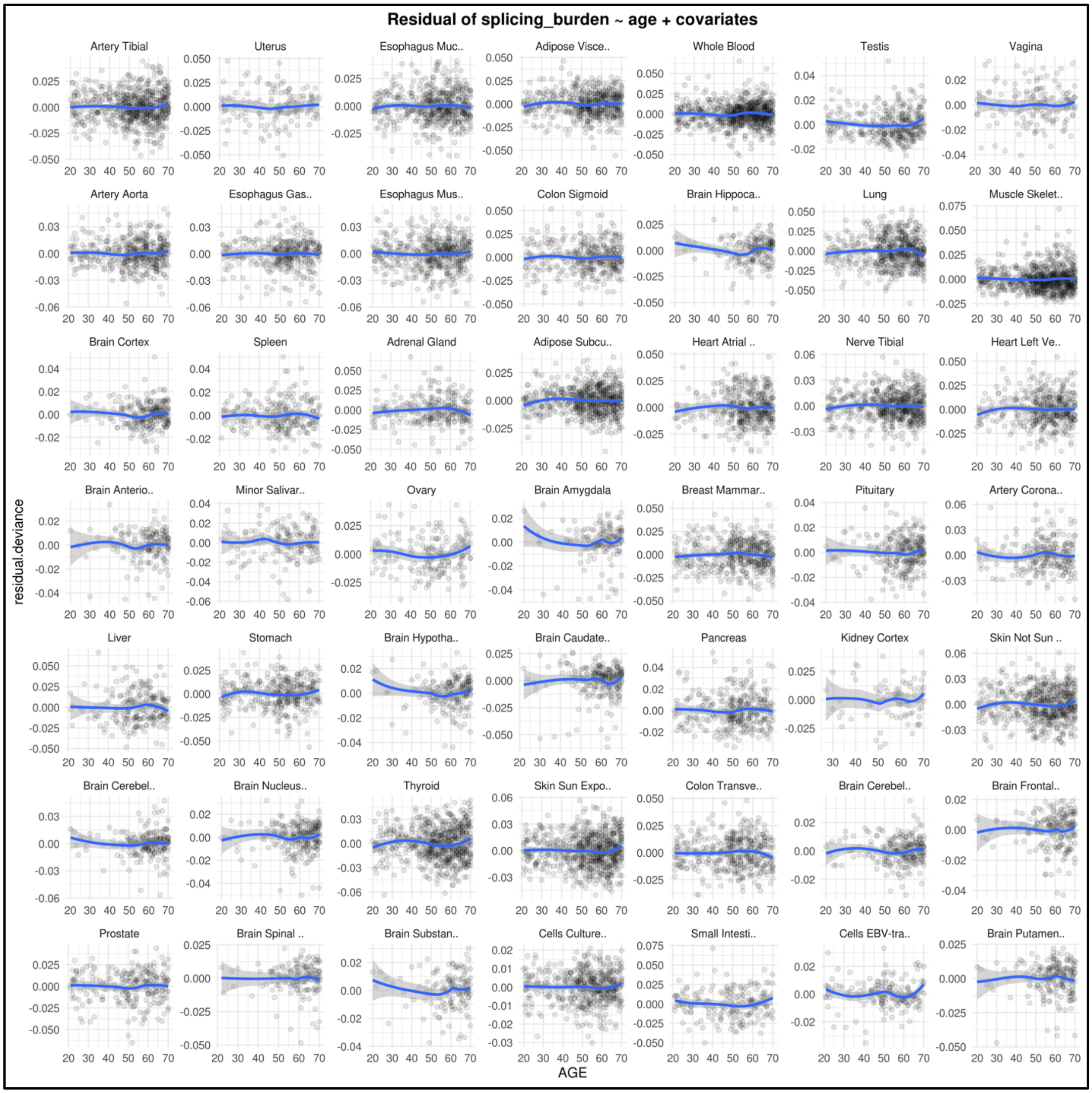


**Fig. S12**. Residual of splicing burden plotted against age. Each panel shows a different subtissue.

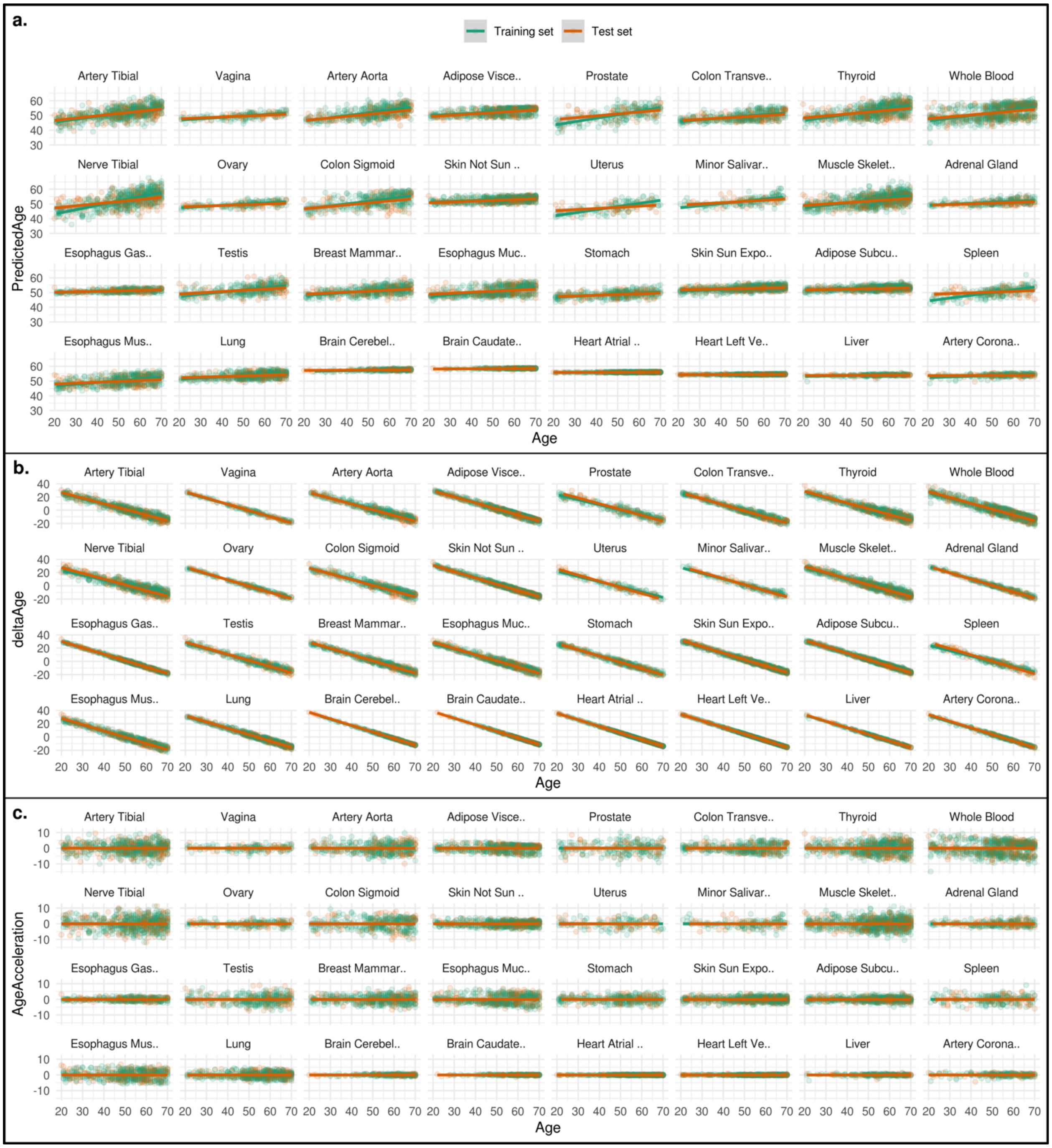


**Fig. S13**. Scatterplots of age predicted by splicing damage clocks. Age is plotted against the age predicted by the given subtissue-clock (panel (a)), or the difference between predicted and real age (delta age; panel (b)), or the residual of predicted age after regressing out age (age acceleration; panel (c)). Subtissues are ordered as in Fig. 7a.

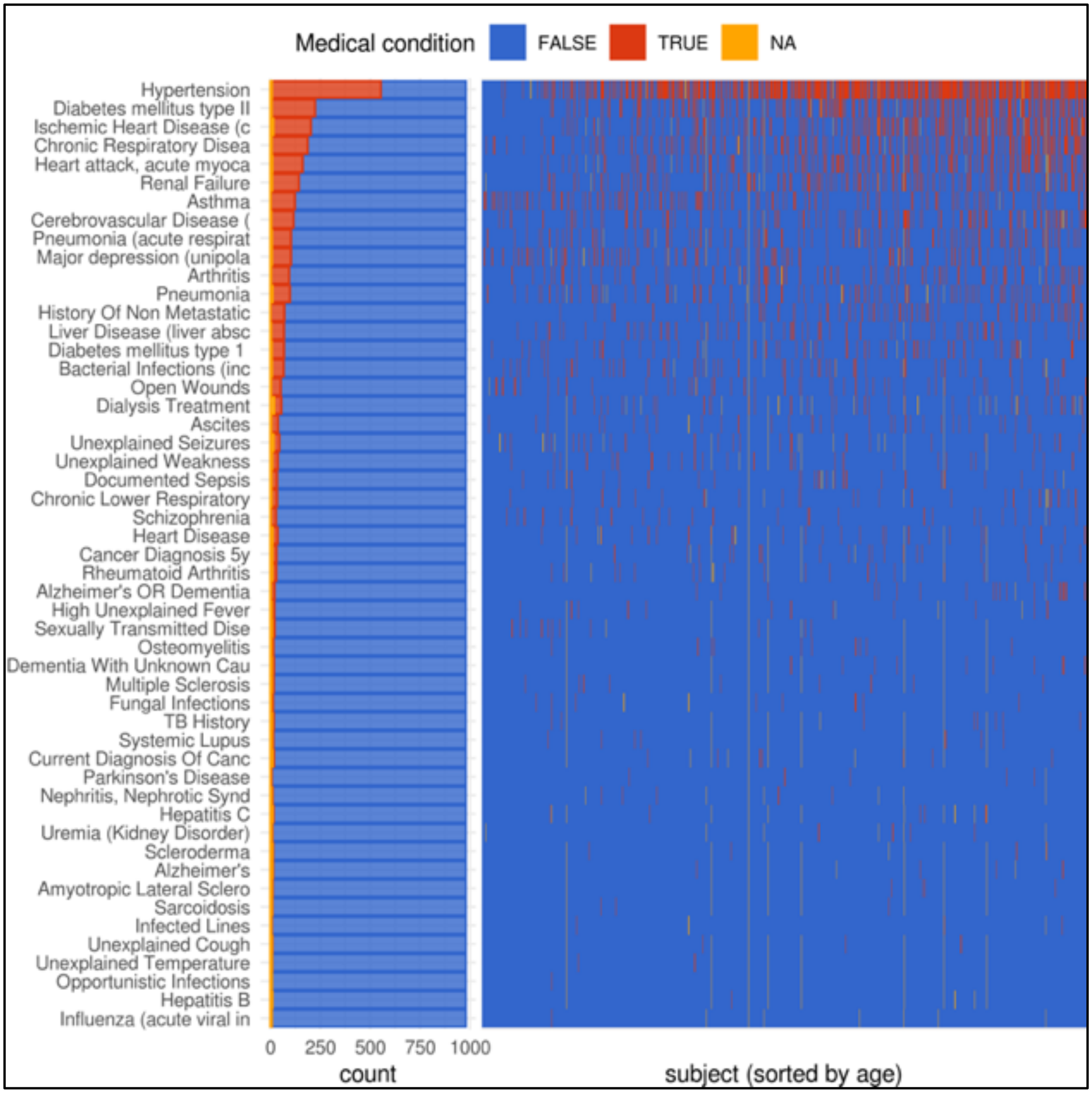


**Fig. S14**. Overview of medical conditions considered in this study. The bar plot on the left shows the occurrence (number of GTEx subjects) of 51 medical attributes that we selected as indicator of disease. The plot on the right shows these annotations in individual subjects (columns), which are sorted by age (the oldest to the right). Red indicates that a condition is present, blue indicates it is not present, and orange indicates that the information is missing.

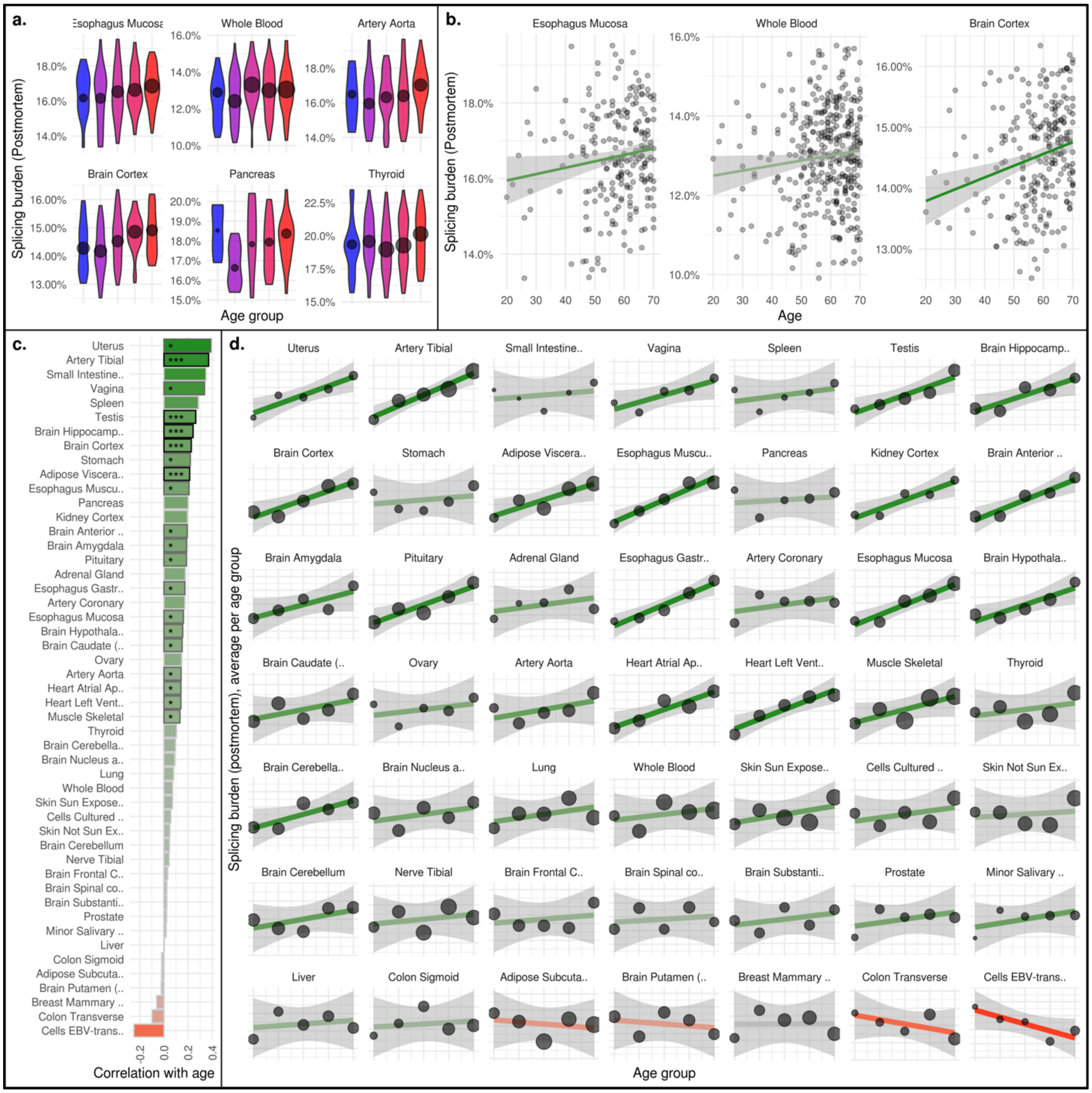


**Fig. S15**. Age changes of mean intron retention in samples from the postmortem cohort. See Fig. 1 for explanation of panels.
